# Rapid and Efficient Quality Control Analysis of Isolated Mitochondria by Interferometric Light Microscopy

**DOI:** 10.1101/2025.05.17.654679

**Authors:** Sabah Mozafari, Christopher Ribes, Dmitry Ayollo, Florence Gazeau, Amanda K. A. Silva, Kelly Aubertin

**Author notes:** Those authors contributed equally to this work (Amanda K. A. Silva, Kelly Aubertin).

## Abstract

Mitochondrial transplantation is a promising biotherapeutic strategy to restore cellular energy homeostasis in diseases associated with mitochondrial dysfunction. However, its clinical translation is hindered by the lack of rapid, reliable, and label-free techniques for quantifying and characterizing freshly isolated mitochondria. Current methods are often time-consuming, costly, or require pre-labeled sample, limiting their suitability for routine quality control in mitochondrial research, engineering, and biotherapy. In this study, we employ interferometric light microscopy (ILM) via Videodrop technology to assess mitochondrial concentration and size with high precision and efficiency. Mitochondria were isolated from human mesenchymal stromal cells (hMSCs) and comprehensively characterized, including through functional and structural marker expression, protein quantification, membrane integrity, and electron microscopy. ILM measurements showed strong correlation and robust accuracy compared to transmission electron microscopy (TEM) and total protein assays. Our findings establish ILM as a label-free, rapid, and scalable approach for mitochondrial quantification and sizing, paving the way for standardized quality control in mitochondrial biotherapy.

## Introduction

Mitochondria are double-membrane organelles found in eukaryotic cells. They play critical roles in maintaining cellular homeostasis, energy production, and metabolism. Mitochondrial dysfunction has been observed in various diseases characterized by metabolic imbalance, including neurodegenerative disorders, cardiovascular conditions, gastrointestinal diseases, and neuromuscular conditions^1–3^. To date, there is a lack of well-established treatment approaches, specifically tackling the prevention or reduction of mitochondrial dysfunction for the improvement of energy metabolism in these diseases.

Physiologically, mitochondria can be transferred to recipient cells via tunneling nanotubes, intercellular dendrites, extracellular vesicle (EV) cargo, or through direct extrusion from donor cells, followed by internalization by recipient cells^4^. Interestingly, studies have shown that blood in a normal physiological state contains a significant number of circulating cell-free, respiratory-competent mitochondria, ranging from 200,000 to 3.7 million per milliliter^5^. Inspired by these natural mechanisms of mitochondrial transfer, exogenous mitochondrial transplantation has been proposed as a potential cell-free biotherapeutic approach for modulating energy metabolism^6^. This concept was first introduced by McCully et al. in 2009 through the local injection of healthy mitochondria into a rabbit model of ischemic heart injury^7^. More recently, mitochondrial transplantation has gained further validation through a 2017 study conducted at Boston Children’s Hospital, which investigated autologous mitochondrial transplantation in the heart as a treatment for myocardial ischemia-reperfusion injury, particularly in pediatric patients^8^.

In recent years, there has been an increasing interest in mitochondria-based biotherapies with research exploring their potential for treating various medical conditions^9^. Currently, more than 10 clinical trials on mitochondrial transplantation are underway worldwide^10^, highlighting the potential of this approach in medical practice. In line with this growing interest, several companies have recently been launched, focusing on mitochondria as a medicinal product, including Cellvie (2018, USA), LUCA Science Inc. (2018, Japan), Mitrix Bio (2018, USA), and MitoSense Inc. (2019, USA). However, several challenges must still be addressed before mitochondrial transplantation can be widely implemented in clinical practice. These include identifying the optimal cell sources, refining technologies for mitochondrial isolation and purification, establishing robust methods for quantification and characterization, developing strategies for long-term storage, optimizing pharmaceutical formulations, and designing effective delivery systems^11,12^.

As a cell source, human mesenchymal stromal cells (hMSCs) offer significant advantages for regenerative medicine due to their accessibility, immunomodulatory properties, and capacity for mitochondrial donation^13^. Additionally, they secrete a diverse array of bioactive molecules, including growth factors, cytokines, and extracellular vesicles (EVs)—collectively known as the secretome—which confer anti-inflammatory, immunomodulatory, and tissue-repairing properties, making them highly effective for various medical conditions^14–18^. hMSCs can be obtained from various sources^19^, including bone marrow, adipose tissue, and umbilical cord, and can be expanded in culture without losing their therapeutic potential. Moreover, their low immunogenicity enables their use in allogeneic settings without provoking strong immune responses^20^. However, compared to traditional cell therapy, which involves administering live cells directly into the patient’s body, cell-free therapy using hMSCs eliminates the risks associated with the engraftment of living cells, such as immune rejection or uncontrolled proliferation. It also provides multifaceted benefits, including tissue repair and immunomodulation, making it a promising and safer alternative to cell-based approaches. For similar reasons, hMSCs are considered one of the best cell sources for cell-free biotherapy based on mitochondria^21,22^, as demonstrated in both preclinical studies and recent clinical trials. This approach represents a potentially revolutionary advancement in regenerative medicine, with a wide range of clinical applications.

Nevertheless, efficiently measuring mitochondrial concentration and size has become a critical challenge for determining mitochondrial characteristics and yield before transplantation. Due to the absence of a simple, rapid, reliable, and universally accepted quantification method, the appropriate dose of mitochondria for transplantation has yet to be standardized. This hampers the ability to compare data across different studies and to establish therapeutic doses, making it difficult to achieve successful mitochondrial transplantation and to translate this approach from bench to bedside. Moreover, due to the instability of extracellular mitochondria—particularly the loss of viability and function following freezing and thawing processes—research in mitochondrial biology and biotherapy has primarily focused on freshly isolated mitochondria. Additionally, most current mitochondrial isolation methods are time-consuming, which can reduce mitochondrial quality and impact transplantation success. Therefore, identifying a rapid and straightforward method for mitochondrial characterization following isolation and prior to transplantation is crucial for ensuring the success and efficacy of mitochondrial transplantation therapies.

So far, various techniques have been developed to determine the concentration and size of isolated mitochondria. For concentration measurement, these methods mainly include the use of a hemocytometer, immunofluorescent staining^7,23,24^, flowcytometry^5,25^, total mitochondrial protein amount^11,26^, or particle counters based on impedance measurements^27^. Some of these techniques, such as flow cytometry and particle counters, can also provide diameter information for isolated mitochondria preparations in addition to counting. However, most of these widely used methods are time-consuming, expensive, imprecise (not automated), or require sample pre-labeling. Techniques like particle counters based on impedance measurement are automated and can provide diameter information, but they require large sample volumes and electrolyte solutions, have no complex workflow, and come with high running costs and a limited size detection range.

In 2016, an analytical device was developed for rapid quantification and sizing of viruses from environmental sources based on interferometric light microscopy (ILM). Since its development, this technic was used for physical characterization of microorganisms in marine water^28^, small cellular particles in natural sources such as blood preparations^29^, viral vectors^30^ (lentivirus and adenovirus), bacteriophages^31,32^ or extracellular vesicles^31,33–35^. The Videodrop is a new, label-free, highly time-saving and cost-effective setup that uses ILM for particle detection and tracking.

In this study, we leveraged Videodrop as a novel and powerful setup to investigate the feasibility of using the ILM technique serving as a rapid and essential quality control analysis for isolated human mitochondria assessing their concentration and diameter. Mitochondria were isolated from hMSCs and underwent comprehensive characterization, encompassing outer membrane marker expression, viability, protein concentration, functional components, membrane integrity, physical and morphological features. Subsequently, the protein concentration of mitochondria was compared to the concentration measured by ILM. Finally, the size characteristics of mitochondria, as determined by TEM, were juxtaposed with the data obtained through ILM. The primary objective was to ascertain whether Videodrop is sensitive enough to precisely and reliably measure the concentration and size of hMSCs-derived mitochondria, serving as an initial quality control step for bioproduction and biotherapy.

## Results

### Isolation of viable and structurally intact mitochondria from hMSCs

Human adipose derived mesenchymal stem cells (MSCs) were successfully expanded in 2D culture flasks. Mitochondria were isolated from hMSCs using the Abcam mitochondria isolation kit in nine independent experiments - each with varying cell numbers (n=9) ranging from 15 to 270 million cells determined by Nucloecounter. The number of cells (in millions) for each experiment is summarized in Table 1. The cell viability measured by Nucleocounter was between 92-97% for all the experiments.

**Table 1.** Mitochondria were isolated from nine experiments with varying cell numbers measured by Nucleocounter, ranging from 15 million cells (Experiment 1) to 270 million cells (Experiment 9), as shown in the table.

| Exp. N° | 1 | 2 | 3 | 4 | 5 | 6 | 7 | 8 | 9 |
| --- | --- | --- | --- | --- | --- | --- | --- | --- | --- |
| Cell number (Millions) | 15 | 30 | 30 | 35 | 35 | 35 | 104 | 115 | 270 |

Isolated mitochondria were characterized in terms of their (i) **viability**, using the MitoTracker Red probe; (ii) **identity**, via immunolabeling of TOM20, a receptor component of the TOM complex (translocase of the outer membrane) in the outer mitochondrial membrane (OMM), as well as Western blotting of mitochondrial oxidative phosphorylation complexes and membrane integrity markers; (iii) **protein concentration** using the Bradford assay; and (iv) **morphology** via transmission electron microscopy (TEM) with negative staining as well as cross-sectioning techniques. Finally, ILM technology was employed using the Videodrop setup to physically characterize the isolated mitochondria in terms of their **quantity** and **size** (hydrodynamic diameter), side by side, compared to the commonly used characterization techniques mentioned above.

MitoTracker Red CMXRos probe was used to stain freshly isolated mitochondria. It is a permeable fluorescent dye containing a mildly thiol-reactive chloromethyl moiety whose accumulation relies on the mitochondrial membrane potential (via the electrostatic interactions of the dye’s positive charge and the negatively charged mitochondrial matrix) ^36,37^. The results confirmed the isolation of numerous viable MitoTracker Red+ mitochondria (Fig. 1a) for all independent experiments (n=9). Counterstaining of samples for the mitochondrial outer membrane marker TOM20 and MitoTracker Red confirmed the presence of multiple MitoTracker Red+/TOM20+ mitochondria (Fig. 1b). Additionally, several TOM20+/MitoTracker-particles were detected following double staining, indicating the presence of non-viable or non-intact mitochondria. This may be attributed to the fact that immunolabeling of mitochondria is a time-consuming process, which could result in the partial loss of mitochondrial integrity.

**Figure 1.**
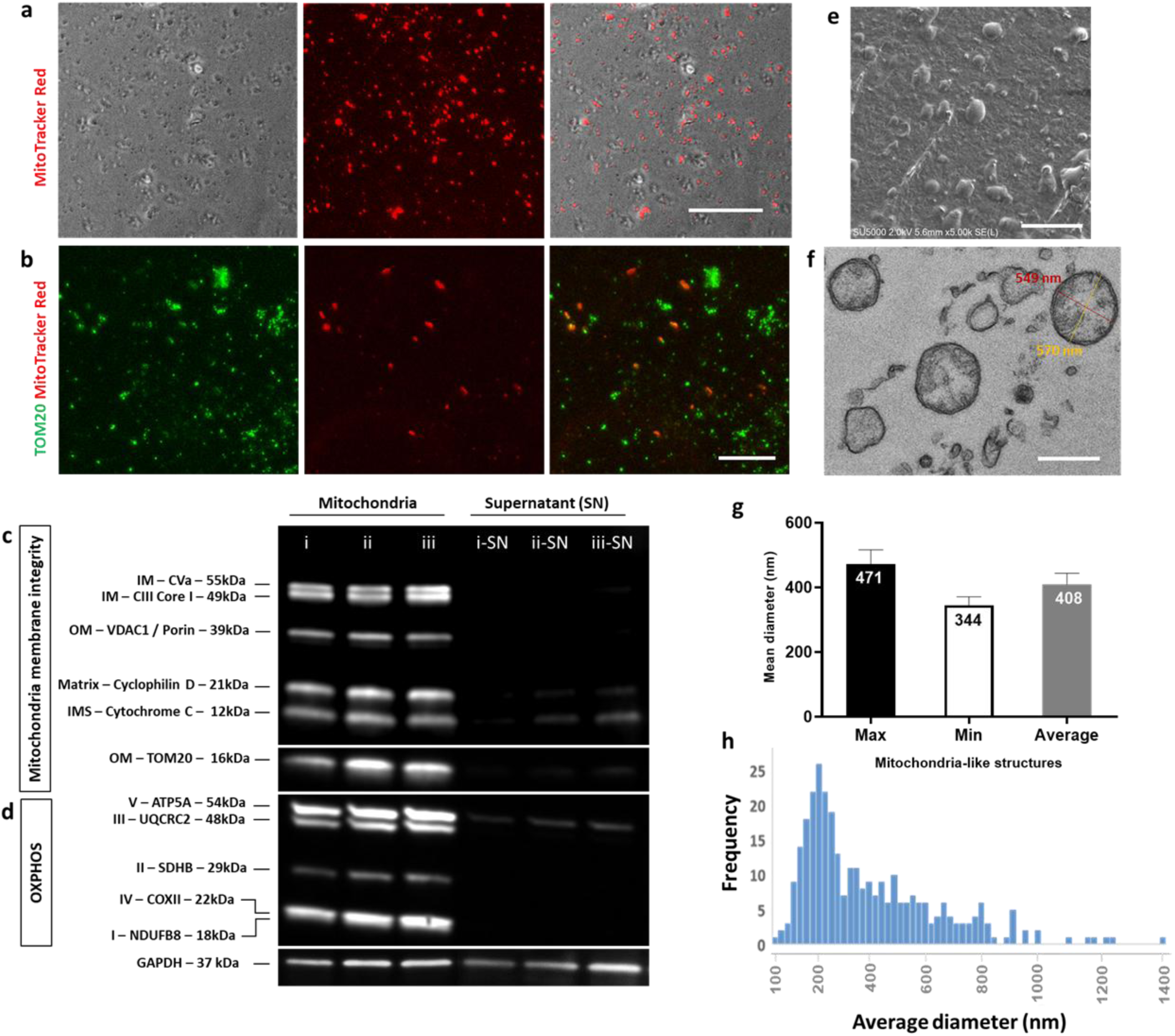
Characterization of Mitochondria Isolated from hMSCs. **(a)** Phase-contrast and MitoTracker Red staining images of isolated mitochondria from hMSCs. **(b)** Fluorescence microscopy images showing numerous TOM20⁺/MitoTracker Red⁺ double-stained mitochondria. Quality control analysis by WB for **(c)** mitochondrial structural and membrane integrity markers as well as **(d)** human OXPHOS complexes (I–V) from three different samples (i-iii). Post-spin supernatants from the same experiments (i–SN, ii–SN, iii–SN) were used as controls. **(e)** SEM imaging revealing multiple mitochondria-like nanostructures (0.2–2 µm in size) with intact outer surfaces. **(f)** Ultrastructural analysis of isolated mitochondria by cross-sectioning TEM, displaying structurally intact mitochondria with maximum (yellow), minimum (red) axis diameters depicted in **(g)**. **(h)** Size distribution graph for mitochondria average diameter measured by TEM. One-way ANOVA followed by Tukey’s multiple comparison test (data taken from n = 4 independent experiments). Error bars represent SEM. SN: supernatant; IM: inner membrane; OM: outer membrane; IMS: intermembrane space; Max: maximum (largest mitochondria length); Min: minimum (shortest mitochondria length); scale bars in a: 50 μm, b: 25 μm, e: 5 µm and f: 500 nm.

The structural integrity of isolated mitochondria was evaluated using Western blotting (WB) analysis (20 µg protein) with the MitoProfile membrane integrity antibody cocktail, which contains 5 antibodies against representative proteins for each of the primary mitochondrial structures: Outer Membrane – Porin; Intermembrane Space – Cytochrome C; Inner Membrane – Complex Vα and Complex III Core 1; Matrix Space – Cyclophilin D. The data show the coexistence of all these structural proteins in the hMSCs-derived mitochondrial preparations (from three different preparations), confirming the isolation of structurally intact mitochondria proteins (Fig. 1c). The presence of TOM20 (another outer membrane marker) was further confirmed by WB of TOM20 protein levels. The post-spin supernatant was used as a control and loaded with the same total protein amount as the mitochondrial samples (20 µg). The results confirmed minimal loss of Cytochrome C, Cyclophilin D, Porin, Complex Vα, Complex III, or TOM20 in the supernatant after the mitochondria isolation process (Fig. 1c, i-SN to iii-SN), therefore validating the isolation method.

Mitochondrial quality was further assessed by WB analysis based on the relative levels of the 5 oxidative phosphorylation (OXPHOS) complexes: Complex I subunit NDUFB8, Complex II subunit 30kDa, Complex III subunit Core 2, Complex IV subunit II, and ATP synthase subunit alpha. The antibodies in the OXPHOS cocktail were selected because they target subunits that are labile when their respective complexes are not assembled. The concurrent presence of total OXPHOS complexes was observed in the isolated mitochondria preparation, confirming the presence of functional mitochondrial elements (Fig. 1d, i-iii shows representative analysis from the triplicate experiments). In contrast, the post-spin supernatant exhibited minimal expression of OXPHOS complexes, confirming the successful isolation of mitochondria (Fig. 1c-d, i-SN to iii-SN). The expression of GAPDH, a housekeeping protein, was detected in both the mitochondria samples and their supernatants. The presence of this cytoplasmic marker alongside 11 structural and functional mitochondrial proteins suggests that the preparations are enriched in mitochondria but not entirely pure.

Morphological and ultrastructural integrity of the isolated mitochondria was further confirmed by scanning electron microscopy (SEM) and transmission electron microscopy (TEM). Indeed, SEM imaging revealed multiple spherical- to oval-shaped structures with intact outer surfaces, ranging in size from 0.2 to 2 µm (Fig. 1e). Furthermore, TEM images demonstrated the presence of intact mitochondria at the ultrastructural level (Fig. 1f). Mitochondria are known for their high plasticity, enabling them to alter their size and shape (ranging from oval to spherical) in response to various cellular conditions and metabolic demands. Additionally, mitochondrial size can vary across different cell types and can be affected by isolation process. To the best of our knowledge, no previous studies have specifically reported the size characteristics of mitochondria isolated from hMSCs. Therefore, the aim was to investigate the size characteristics of isolated mitochondria derived from hMSCs.

To achieve this, three key parameters were measured for each mitochondria-like structure observed in the TEM images of cross-sectioned samples images: the maximum axis diameter (Fig. 1f, shown in yellow), the minimum axis diameter (Fig. 1f, shown in red), and the average of these two diameters. A total of 416 mitochondria-like structures were analyzed across four independent experiments (n = 4), with 3–5 measurement fields per experiment at ×25k magnification.

The results showed that the maximum size ranged from 116 to 1700 nm, with a mean maximum diameter of 471.9 ± 44 nm. The minimum diameter ranged from 92 to 1049 nm, with a mean minimum diameter of 344.4 ± 26 nm. The overall average diameter ranged from 105 to 1346 nm, with a mean value of 408.2 ± 35 nm (Fig. 1g). These values are summarized in Supplementary Table 1.

Statistical analysis revealed no significant difference between the maximum, minimum or the average diameters of the isolated mitochondria-like structures. However, the mitochondria-like structures were found to have conserved partially the anisotropic geometry after isolation as illustrated by the difference between max and min diameters. The average size distribution of isolated mitochondria-like structures is depicted in Fig. 1h showed that hMSC-derived and isolated mitochondria are highly polydisperse. At the ultrastructural level, approximately 5.29%, 0.72%, and 1.92% of all quantified mitochondria-like structures had diameters exceeding 1 µm based on maximum, minimum, and average diameter measurements, respectively. Supplementary Table 1 summarizes the size characteristics of the isolated mitochondria-like structures.

### Mitochondria concentration measurement by ILM

The Videodrop instrument (Myriade, France) is a label-free, rapid, and cost-effective method that allows sub-micrometric particle detection and tracking (Fig. 2). It is able to characterize the size (hydrodynamic diameter) and the concentration of (bio)nanoparticles using only few microliters (5 to 10µL at ∼10^9^ part/mL) of the precious samples in less than a minute per sample with the mains advantages of being easy-to-use, fast and in a non-denaturant manner and operated without any fluidics (reduced probability of bubbles, no clogging). The system does not require prior calibration nor maintenance from the user. Additionally, it can be used to characterize nano-objects in viscous or complex fluids such as pharmaceutical formulation^38^.

**Figure 2.**
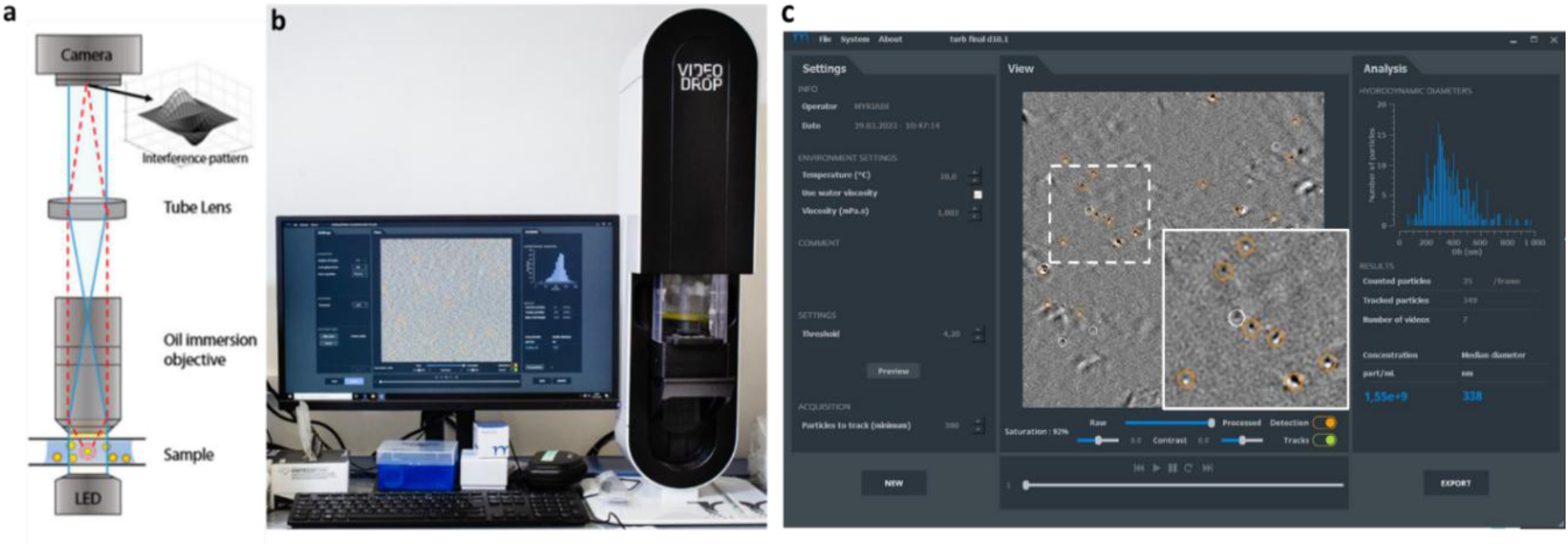
Particle size and concentration characterization by ILM using Videodrop: (a) Schematic diagram of the optical setup. Videodrop technology is based on ILM that uses a LED light source to illuminate the sample. Its optical system consists of a transmission brightfield microscope equipped with an oil immersion objective and a camera collecting images. The interference between the incident light and light scattered by all particles creates an interference pattern. The interference images are obtained by processing the microscope images by a QVIR software and allow to detect the particles; (b) Photograph of the Videodrop system connected to a computer equipped with QVIR software; (c) Screenshot displaying the QVIR software during a measurement. The inset image provides a closer view of the interferometry signals in a white dotted box, on which we can visualize particles. The detected particles (not tracked) are displayed in white and the detected and tracked particles are displayed in orange. The hydrodynamic diameter (size) is calculated from the diffusion coefficients of tracked particles only. All detected particles (white + orange) contribute to the number of particles per volume unit (concentration).

The Videodrop is a transmission brightfield microscope employed as an interferometric light microscope that allows the visualization of sub-micrometric particle from interference patterns. The system contains a camera capturing videos, and is associated with an image processing algorithm able to detect and track nanoparticles^39,40^. Figure 2 shows the Videodrop instrument, its optical set-up and depicts the principle of the measurement based on ILM (Fig. 2). Counting particles allows to measure the concentration, while tracking their Brownian motion enables to measure individually their hydrodynamical diameter by using the Stokes–Einstein equation (Eq. 2) and to provide size distribution by plotting the number of particles for hydrodynamic diameter ranging from 50 to 1000 nm (Fig. 2c). Fig. 2b-c show the software measurement views.

The interferometry-based analytical method was utilized to simultaneously quantify the concentration and hydrodynamic size distribution of hMSC-derived mitochondria using Videodrop. First, we focused on quantifying mitochondria concentration. Isolated mitochondria samples from the 9 independent experiments summarized in Table 1 were analyzed separately using Videodrop.

The sensitivity and accuracy of the Videodrop for concentration and diameter assessment depend on relative positioning of the sample concentration regarding the optimal range of the instrument. Indeed, an over-estimation of the size can be observed when the sample concentration is too high. For nanoparticles such as extracellular vesicles, this range is known to be between 10^8^ and 10^10^ (centered around 1*10^9^) part/mL. In order to estimate the concentration windows that gives the most reliable Videodrop measurements for mitochondria concentration (ie the linear range for the concentration of mitochondria), measurements were done at 5-7 serial dilutions for each mitochondria preparation (N=9). Each Videodrop measurement was performed on triplicate samples with a set of minimum 300 particles to be tracked. Fig. 3a-f depicts representative interferometry images and the corresponding size distribution of an experiment in which 6 serial dilutions (serial dilution factors 1, 2, 4, 8, 16 and 32, ie dilution factors of d50, d100, d200, d400, d800, and d1600 from the isolated mitochondria samples) were analyzed by Videodrop. As expected, interferometry images visibly show a decrease in the particle density while the dilution factor increases. It is observed that the mean diameter of particles seems to converge to a value while the dilution factor increases. The particle mean size influences the size distribution of the particles, with higher concentrations exhibiting larger particle sizes possibly due to particle aggregation or because mitochondria interact with each other mechanically reducing their movement (sub-diffusion) and inducing therefore an under-estimation of their size. Indeed, at the highest dilution factor, the mean size was lower (277 ± 2 nm), while at the lowest dilution factors, it was higher (441 ± 4 nm). At low concentrations, limited particle sampling can reduce measurement reliability.

**Figure 3.**
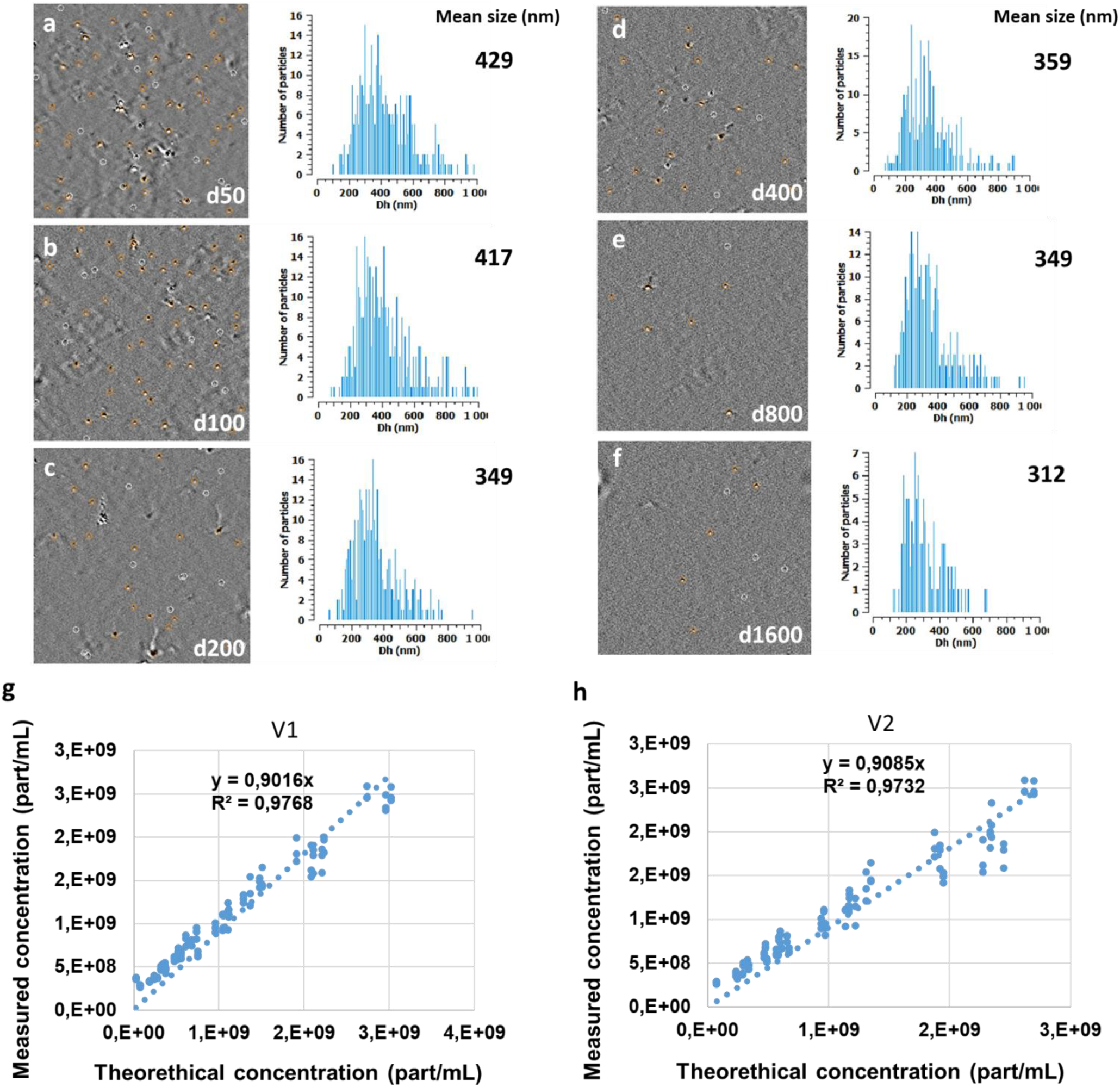
Characterization of Isolated Mitochondria Using Interferometry Light Microscopy with Videodrop. Mitochondrial preparations from 9 independent experiments were physically characterized using Videodrop, with multiple serial dilutions performed in triplicate. **(a-f)** Representative ILM Images from one experiment, along with corresponding size distribution graphs and measured mean sizes, are shown for serial dilution factors of d50, d100, d200, d400, d800, and d1600. A gradual decrease in particle density and a reduction in mean particle size are observed as the dilution factor increases, ranging from 429 nm (dilution factor 1) to 312 nm (dilution factor 32). **(g-h)** Data from all nine experiments (5–10 serial dilutions, each performed in triplicate) demonstrate linearity within an optimal concentration range of 7.5 × 10⁷ particles/mL to 2.7 × 10⁹ particles/mL (V1) or 9.5 × 10⁸ to 2.5 × 10⁹ particles/mL (V2) for mitochondria-rich samples.

Identifying the accurate windows for mitochondria concentration measurement is equivalent to identifying the range for which the measured concentration is linear (proportional) with the theoretical concentration (and inversely proportional to the dilution factor). As is it not possible to retrieve the real theoretical concentration, we defined as the theoretical concentration the value that should be the closest for the real number of particles per unit of volume. We developed two different methods to estimate the “theoretical” concentration of each sample based on the combination of measurements performed at different dilution factors, by either (method V1) identifying the concentration that was the closest to 1*10^9^ part/mL, which is the center of the optimal working range for concentration quantification using Videodrop, or (method V2) to average all the concentrations that are given a director coefficient close to 1 when fitting the plot of log(measured concentration) as a function of log(1/dilution factor) (Fig3.g-h). A schematic of the method V2 can be found in supplementary figure 1.

Once the theoretical concentration was found for each of the 9 independent experiments individually, we plotted the measured concentration as a function of the theoretical concentration for the 9 experiments on the same graph (Fig. 3-g and h for method V1 and V2, respectively). First, we can observe for both methods a good linearity between the theoretical and the measured concentration. Second, while method 2 is more robust because of the consideration of several measurements (performed at different dilution factors), the results obtained were of similar efficiency with both methods. Indeed, by comparing the linear fitting performance (ie R^2^ coefficient values) of Fig. 3g and 3h, we can see that both methods gave similar fitting coefficient and similar R^2^ coefficients.

Using these complementary methods, the linear concentration range for Videodrop was established, highlighting the particle density range where the system accurately measures mitochondrial particle concentrations across different experiments. Within this range, particle numbers measured by Videodrop varied linearly with particle density, ensuring reliable and consistent quantification.

Moreover, our results indicate that the optimal density of interferometry signals for mitochondria-rich preparations falls within the range of 20-30 orange and 5-10 white signals per frame (totaling 30-40 particles). This corresponds to a measured concentration range of 9.5 × 10⁸ to 2.5 × 10⁹ particles/mL (intersected range of V1 and V2 methods), based on tracking a minimum of 300 particles.

The particle size distribution obtained from individual particle tracking (Fig. 3a-f) showed consistent profiles within the above determined linear concentration range.

Interestingly, thanks to ILM quantification, we observed a good correlation between the particle quantity and the cell quantity from which mitochondria were isolated. The total particle counts, as measured by Videodrop for the nine experiments with varying cell numbers (Table 1), are illustrated in Fig. 4a. A linear correlation was found between the total particle count measured by Videodrop and the total cell number (measured by NucleoCounter®) originally used for the production of these particles, with an *R*^2^ value of 0.9649 (Fig. 4b). Videodrop data analysis revealed a mean production of 3513 ± 148 particles per cell.

**Figure 4.**
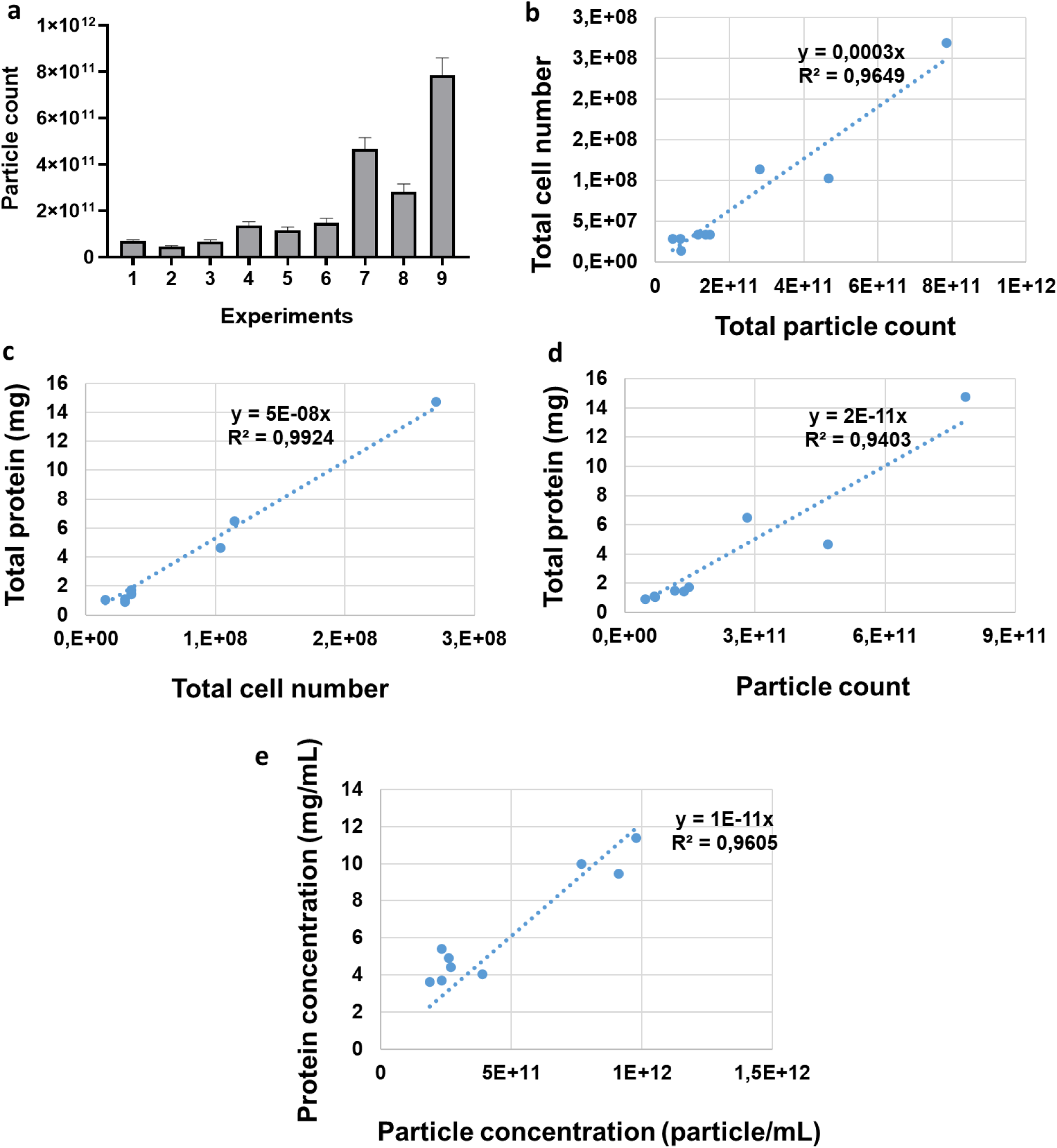
Correlation Between Mitochondria Concentration Measured by Videodrop and Protein Concentration. **(a)** Total particle count measured by Videodrop across 9 independent experiments. **(b)** Videodrop particle count correlates with the total cell number determined by NucleoCounter **(c)**. A strong correlation is observed between the total cell number and the total protein amount (mg) measured by the Bradford assay from the mitochondria extract. **(d)** Additionally, a robust correlation is found between the total protein amount (mg) and the total particle count measured by Videodrop. **(e)** A similar correlation is observed between protein concentration (mg/mL) and Videodrop concentration (particles/mL) in mitochondria samples. n = 9 independent experiments.

To further validate the reliability of mitochondria concentration measured by Videodrop, protein concentration of mitochondria sample (n=9) was measured by Bradford protein assay. Protein concentration measurement is a widely used method to determine the mitochondria concentration^11^. First, a correlation was observed between the total amount of protein in each isolated mitochondria sample and the total number of initial cells with *R*^2^ value of 0.9924 (Fig. 4c). Between 30 and 70 µg of protein (or a mean of 47 ± 4 µg) protein equal to 3.30e+9 ± 3.62e+8 particles) were generated per 1 million hMSCs. The resulting data showed also a strong correlation between the total amount of protein (or the protein concentration) measured by Bradford test and the total particle count (or particle concentration, respectively) measured by Videodrop with *R-squared* values of 0.9403 and 0.9605 for total amount of protein (mg) vs particle counts or protein concentration (mg/mL) vs particle concentration (Particle/mL), respectively (Fig. 4d-e). Consequently, we could determine a mean protein amount of 15.3 ± 1.5 µg for 10^9^ particles counted by Videodrop.

### ILM provides reliable mitochondria size characteristics

ILM analysis using Videodrop can also provide size measurements. Indeed, Videodrop allows to follow the particles in time. From the 2D coordinates of each particle, the mean square displacement (MSD) is calculated. For particles moving freely in a purely viscous liquid, ie particles submitted to Brownian motion, the MSD is proportional to the diffusion coefficient multiplied by the time (Eq. 1). Therefore, a linear fit of the MSD as a function of time allows to quantified the diffusion coefficient of each particle. The size of each particle is then retrieved from the Stokes-Einstein equation (Eq. 2) linking the diffusion coefficient with the temperature, the fluid viscosity and the hydrodynamic radius (valid for spherical particles). More detailed information about ILM size quantification can be found in the paper of Alexandre et al.^35^.

While mitochondria are not necessarily all perfectly spherical, we made the hypothesis that the Stokes-Einstein equation was still valid to evaluate the capability of the Videodrop instrument to quantify properly mitochondria size: we compared the measurements of the hydrodynamic diameter by ILM to the size of isolated mitochondria using TEM techniques of negatively stained vs cross-sectioned resin-embedded samples (Fig. 5).

**Figure 5.**
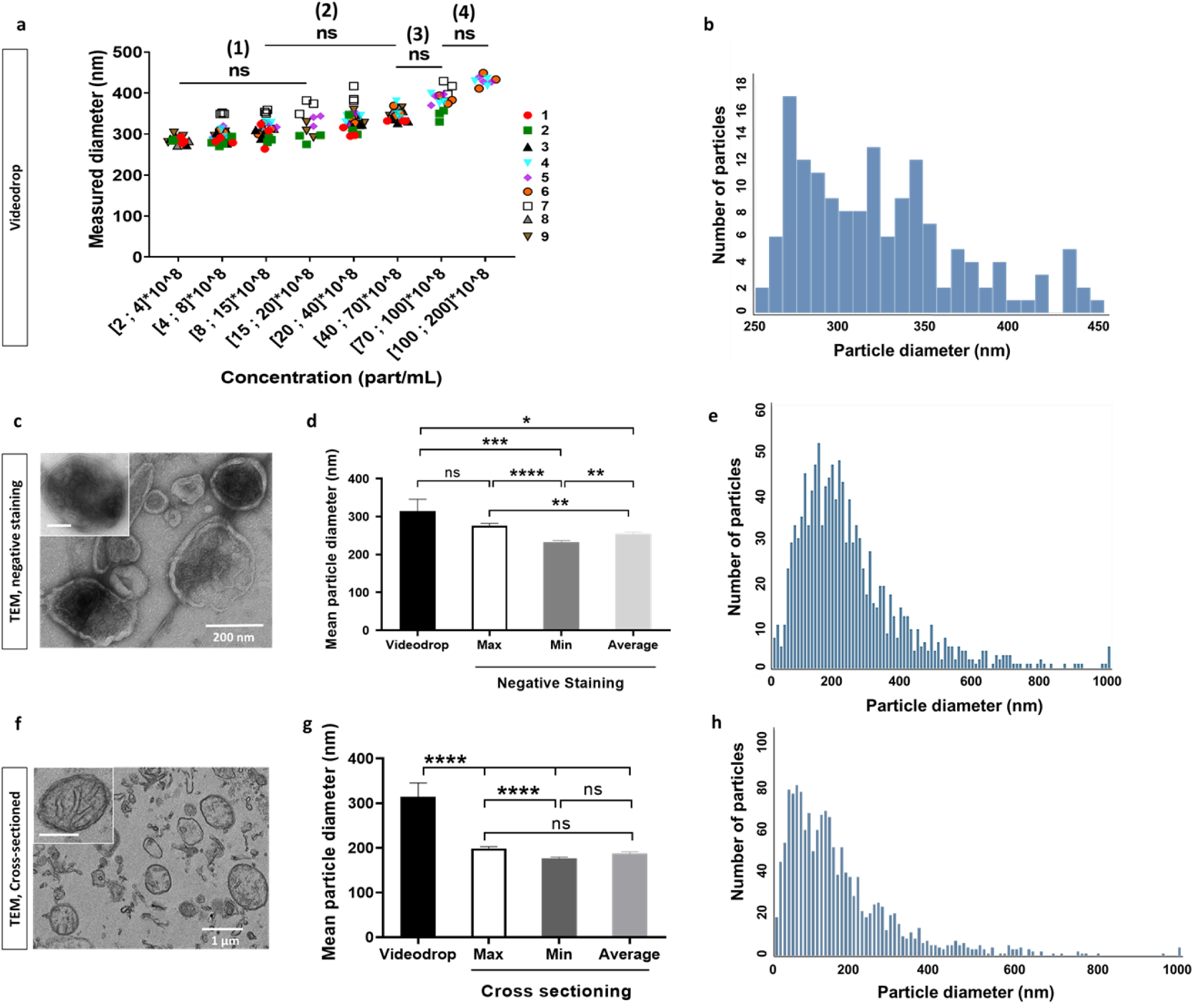
Measurement of Isolated Mitochondria Size Using ILM via Videodrop. **(a)** Determination of concentration domains in which Videodrop provides consistent size measurements for isolated mitochondria samples (n = 9 independent experiments, mean diameter: 315 nm). Four different regions were identified: (1), (2), (3) and (4). **(b)** Size distribution graph for all particles within the linear concentration range shown in (a). **(c)** Representative TEM images of negatively stained mitochondria, revealing multiple double-membrane mitochondria-like structures of various dimensions. **(d)** Mean particle size measured by Videodrop (313 nm) is consistent with the mitochondria maximum diameter (297 nm) but differs from the minimum (229 nm) and average (260 nm) diameters measured by TEM with negative staining (n = 7 experiments). **(e)** Size distribution graph of mitochondria average diameters measured by TEM with negative staining. **(f)** Representative high-resolution TEM images of cross-sectioned mitochondria, confirming intact inner and outer membranes, along with smaller structures of various shapes produced during mitochondria isolation. **(g)** Mean size measured by Videodrop is significantly higher than the maximum (218 nm), minimum (197 nm), and average (212 nm) diameters measured by cross-sectioned TEM (n = 4 experiments). **(h)** Size distribution graph of mitochondria average diameters measured by cross-sectioned TEM. One-way ANOVA followed by the Tukey’s multiple comparison test. * *p < 0.05*, ** *p < 0.01*, *\*\*\* p < 0.001, **** p < 0.0001*. Error bars represent SEM. ns: not significant; Max: maximum, for the largest particle diameter; Min: minimum, for the shortest particle diameter; scale bars in (c): 200 nm, (f): 1 µm, insets in (c and f): 200 nm.

Videodrop mean sizes from all nine experiments (consisting of 5-7 serial dilutions, with triplicates for each dilution) were plotted based on 8 different particle concentration intervals: [2 to 4] × 10^8^, [4 to 8] × 10^8^, [8 to 15] × 10^8^, [15 to 20] × 10^8^, [20 to 40] × 10^8^, [40 to 70] × 10^8^, [40 to 70] × 10^8^, and [70 to 100] × 10^8^ part/mL (Fig. 5a-b).

As mentioned in previous section, the mean size of particles, measured by Videodrop, tends to decrease when the sample concentration is decreased over the full range of concentration, from 429 ± 1 nm to 285.6 ± 1.81 (average over the 9 independent experiments). In order to identified the intervals for which the hydrodynamic diameter was variating non-significantly, we performed statistical analysis be regrouping the above concentrations intervals. We identified 4 concentration regions ((1), (2, (3), (4), Fig. 5a). In the above section, we found that the linear range for concentration is between 9.5*10^8 and 2.5*10^9 part/mL. The concentration regions that matches he best the linear range is the region (2). The size distribution corresponding to this region (2) (Fig. 5b) gave an average value of 315 ± 6 nm (n=9 experiments) for the isolated mitochondria.

Additionally, mitochondria size was assessed by two TEM techniques having different resolutions: TEM with negative staining, which typically provides lower resolution; and TEM with cross-sectioning, known for its high imaging resolution.

First, the TEM images of negative staining were analyzed. Fig. 5c displays intact double membrane mitochondria with spherical to tubular structures, each exhibiting different sizes. For each structure, the minimum, maximum and average diameters were compared to the mitochondria mean diameter measured by Videodrop (Fig. 5d). Figure 5e shows the average size distribution graph for all detected particles by negative staining TEM method (n=7 experiments, 1,214 particles analyzed). Data analysis revealed that across all observed particles, the average size measured by negative staining ranged between 27-1564 nm, with a mean diameter of 254 ± 4.7 nm (Fig. 5c-e). Significant differences were observed between the mean minimum (232 ± 4.4 nm) and maximum (276.2 ± 5.8 nm, *p-value <0.0001*) diameters as well as between both of these values and the average particle diameter (*p-value = 0.0095*), highlighting the anisotropy in mitochondria geometry. Interestingly, while statistically significant differences were found between the mean minimum (*p-value = 0.0002*) or average (*p-value =* 0.014) diameters, no significant difference was detected between the mean maximum and the mean particle diameters measured by Videodrop (313.9 ± 3.5 nm, Fig. 5d). The size distribution for the average diameters measured by negative staining TEM is shown in Figure 5e.

Subsequently, TEM images obtained from cross-sectioning method were analyzed offering a higher resolution compared to negative staining or interferometry images. Figure 5f shows TEM images of cross-sectioned samples from mitochondria-enriched preparations. These images revealed mitochondria of varying diameters, as well as smaller structures such as EVs. Cross-sectioning TEM analysis was performed on a total of 1,503 particles (n=4 independent experiments), revealing an average size distribution ranging from 31 to 1631 nm, with an average diameter of 187.4 ± 3.5 nm. While no significant differences were detected between the average and mean minimum (176.2 ± 3.3 nm) or maximum (198.7 ± 4 nm) diameters, a statistically significant difference was found between the mean minimum particle size and the mean maximum size (*p-value < 0.0001*, all particles from 4 experiments, Fig. 5g), confirming the anisotropic geometry observed with negative staining TEM. However, statistically significant differences were found between all these diameters measured by cross sectioning TEM with the mean diameter measured by ILM using Videodrop (*p < 0.0001,* Fig. 5g). The size distribution for the average diameters measured by cross sectioning TEM is shown in Figure 5h.

As previously mentioned, the optimal size detection range of Videodrop is limited to diameters between 80 and 1000 nm. Analysis of all particles detected by TEM revealed that 5.52% and 17.89% of particles had an average size below 80 nm, while 0.41% and 0.26% exceeded 1000 nm, as measured by the negative staining and cross-sectioning TEM methods, respectively. The majority of particles—94.15% and 81.83%—fell within the 80–1000 nm range for the negative staining and cross-sectioning TEM methods, respectively. Notably, this range accounted for up to 98% of mitochondria-like structures identified using cross-sectioning TEM. These findings are summarized in Table 2.

**Table 2.** Size measurements of mitochondria-like structures and all detected objects by TEM, assessed via cross-sectioning and negative staining methods. Data represent the total number of particles analyzed per condition, collected from n=4 independent experiments for the cross-sectioning method and n=7 independent experiments for the negative staining method.

|  |  |  |  | Range/Mean (nm) |  |  |  |  |  |  |
| --- | --- | --- | --- | --- | --- | --- | --- | --- | --- | --- |
| TEM method | Analyzed particles | Total particles | Diameter | All particles |  | 80 <> 1000 |  | % 80 <> 1000 | % <80 | % >1000 |
| Cross-sectioning | Mito-like structures | 416 | Min | 92 - 1049 | 354.4 ± 8.9 | 92 - 948 | 352.54 | 99.27 | 0 | 5.29 |
|  |  |  | Max | 116 - 1700 | 489 ± 13 | 116 - 978 | 453.73 | 94.71 | 0 | 0.72 |
|  |  |  | Average | 105 - 1346 | 421.7 ± 10 | 105 - 964 | 410 | 98.07 | 0 | 1.92 |
|  | All objects | 1503 | Min | 31 - 1230 | 176.2 ± 3.3 | 80 - 839 | 203 | 79.24 | 20.55 | 0.2 |
|  |  |  | Max | 31 - 2031 | 198.7 ± 4 | 80 - 950 | 223 | 82.1 | 17.56 | 0.33 |
|  |  |  | Average | 31 - 1631 | 187.4 ± 3.5 | 80 - 964 | 211 | 81.83 | 17.89 | 0.26 |
| Negative staining | All objects | 1214 | Min | 27 - 1406 | 232 ± 4.4 | 80 - 997 | 241 | 93.32 | 6.5 | 0.24 |
|  |  |  | Max | 27 - 2734 | 276.2 ± 5.8 | 80 - 997 | 278.9 | 93.73 | 5.51 | 0.82 |
|  |  |  | Average | 27 - 1564 | 254 ± 4.7 | 80 - 997 | 260.8 | 94.15 | 5.52 | 0.41 |

To ensure a more direct comparison between Videodrop and TEM data, the TEM dataset was re-analyzed, considering only objects with a mean diameter greater than 80 nm, as particles exceeding 1000 nm were already negligible in both TEM methods.

The data from negative staining TEM revealed a statistically significant difference between the mean maximum (297.6 ± 16 nm) and minimum (229.7 ± 19 nm) particle diameters (*p = 0.02*). However, no significant difference was detected between the average diameter (260.8 ± 14 nm) and either the maximum or minimum diameters, underscoring the anisotropic shape of mitochondria (Fig. 6a). Furthermore, while the mean diameter measured by Videodrop (315.3 ± 9.8 nm) was significantly larger than the mean minimum diameter measured by negative staining TEM (*p = 0.0035*), no statistically significant differences were observed between the mean size measured by Videodrop and the maximum or average particle diameters measured by TEM (Fig. 6a-b, n = 7 experiments, 4–10 measurement fields at ×70k magnification each).

**Figure 6.**
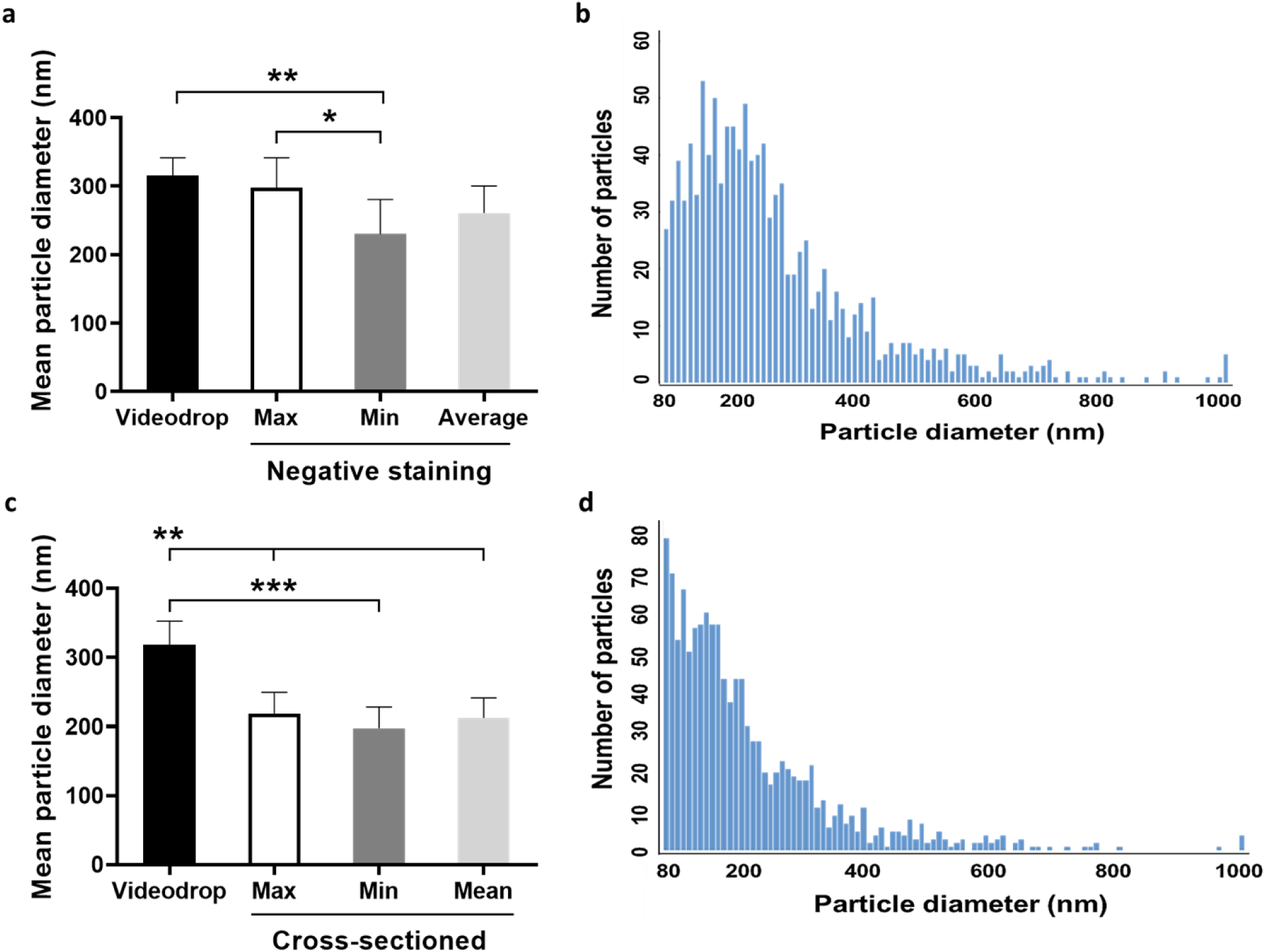
Particle size measurement using negative staining and cross-sectioning TEM techniques (for particles >80nm). **(a)** Data indicating that the Videodrop mean particle size (318 nm) closely aligns with the mitochondria mean maximum (297 nm) and average (260 nm) diameters but not with the minimum diameter (229 nm) measured by TEM using the negative staining method for particles exceeding 80 nm. **(b)** Size distribution of the average particle diameters of isolated mitochondria samples (n = 7 experiments) measured using the negative staining method. **(c)** Comparison of the mean diameters of isolated mitochondria samples measured using TEM cross-sectioning (minimum = 197 nm; maximum = 218 nm; average = 212 nm; n = 4 experiments) with the mean particle diameter measured by Videodrop (318 nm). **(d)** Size distribution of the average particle diameters measured using the cross-sectioning method. Only particles exceeding 80 nm (Videodrop detection limit) were considered for both TEM methods. One-way ANOVA followed by Tukey’s multiple comparison test (b and d). * p < 0.05, ** p < 0.01, *** p < 0.001. Error bars represent SEMs. Max: maximum, for the largest particle diameter; Min: minimum, for the shortest particle diameter.

Once again, the minimum, maximum, and average diameters of particles exceeding 80 nm in TEM images of cross-sectioned samples were analyzed. Statistical analysis showed no significant differences between the mean minimum (197 ± 15 nm), mean maximum (218 ± 15 nm), and mean average (212 ± 14 nm) diameters (Fig. 6c). However, the mean particle size measured by Videodrop (318 ± 17 nm, n = 4 experiments, 12–30 measurements per experiment) was significantly larger than the mean minimum (*p-value = 0.0008*), maximum (*p-value = 0.0036*), and average (*p-value = 0.0023*) diameters measured by cross-sectioning TEM (Fig. 6c-d, n = 4 experiments, 3–7 measurement fields per experiment at ×10k magnification).

### Mitochondria characterization by immunofluorescence microscopy in comparison to ILM

Since mitochondrial staining (either through flow cytometry^5,41^ or fluorescence microscopy^7,23^) is widely used to characterize mitochondria’s physical properties, such as size and density, we employed this approach to compare mitochondrial size and quantification with results obtained using Videodrop and TEM. To minimize quantification bias, automated fluorescence microscopy-based analysis was used. As mentioned above, TOM20, a component of the outer-membrane translocator TOM40 complex, is a mitochondria-specific protein that has been widely used as a mitochondrial marker^42,43^. We used the Cell Insight CX7 LZR confocal spinning-disk microscope, equipped with the automated High-Content Screening (HCS) Studio Cell Analysis software platform (ThermoFisher Scientific), to visualize TOM20+ structures in different mitochondrial preparations (Fig. 7a-b, n = 3 independent experiments). We then automatically quantified their size and density across 10 different fields, with triplicate coverslips per experiment. Blue circles indicate TOM20+ white objects (Fig. 7bi). Typically, numerous aggregated and overlapping TOM20+ mitochondria were observed following immunofluorescence (IF) staining (Fig. 7b). The results revealed a particle size range of 249 nm to 11,717 nm, with a mean size of 938.5 ± 1 nm (total quantified objects: 130,519). The particle size distribution graph for TOM20 IF automated quantification is shown in Fig. 7c.

**Figure 7.**
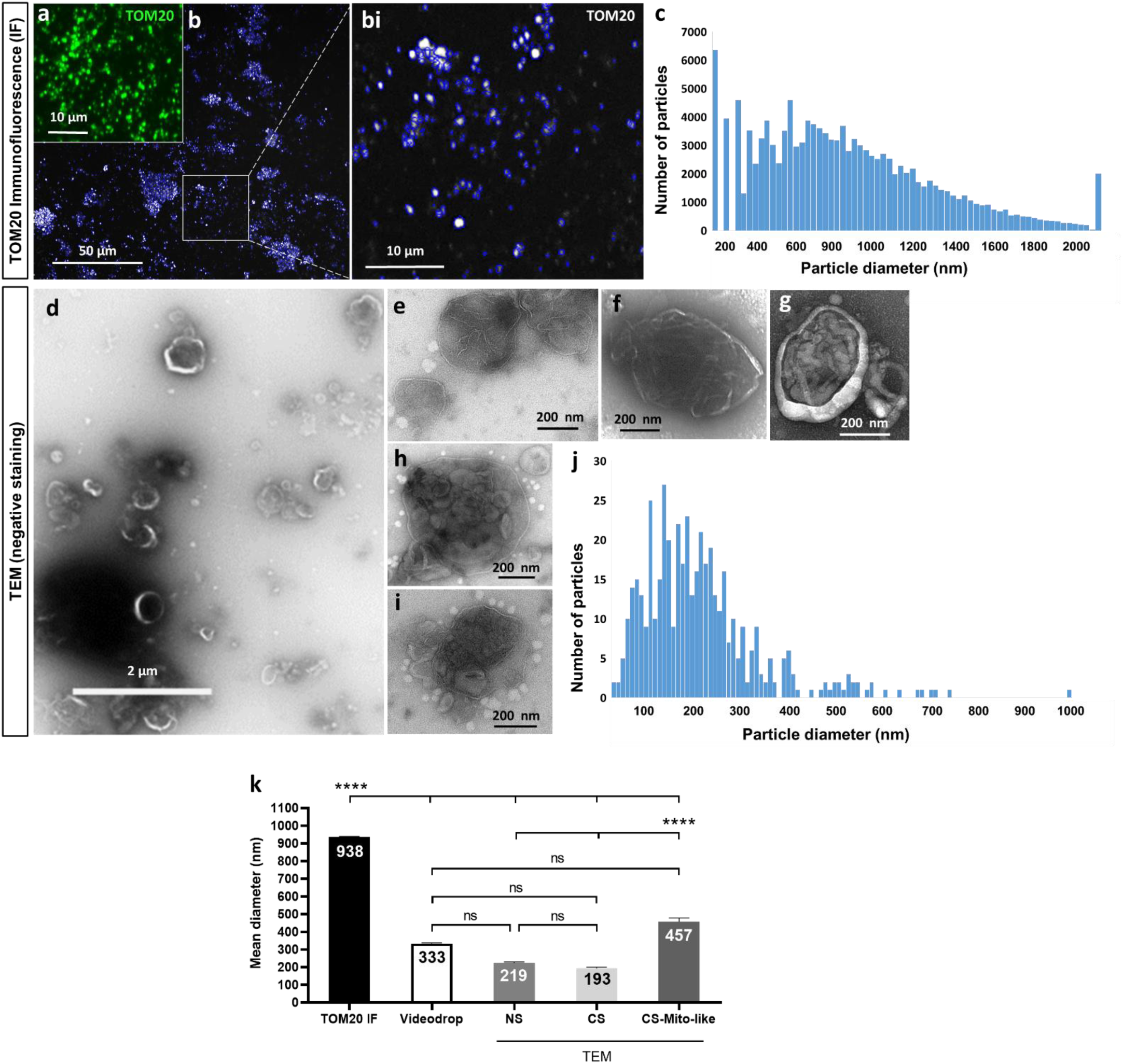
Characterization of isolated mitochondria by IF microscopy vs ILM. **(a)** (a) CellInsight™ confocal microscopy image showing fluorescence labeling of isolated mitochondria with TOM20. (b) Quantification of TOM20+ particles (white spots circled in blue, with a higher-resolution view shown in the inset-bi) measured using automated HCS Studio Cell Analysis software. (c) Particle-size distribution of TOM20+ mitochondria preparations. (d-i) Representative TEM images (negative staining) showing an overview and higher magnification of several mitochondria-like structures (e–i). (j) Particle-size distribution for the same samples detected by TEM (negative staining). (k) Graph showing a significantly larger mean diameter measured by IF or for mitochondria-like structures measured by cross-sectioning TEM. No significant differences were found between the mean particle diameters detected by Videodrop or TEM (negative staining or cross-sectioning methods), including all particles or excluding those with diameters outside the detection range for Videodrop (Supplementary Fig. 1). The same experiments (n = 3) were used for different quantification methods in this figure. Statistical analysis was performed using one-way ANOVA followed by Tukey’s multiple comparison tests (k and l). **** p < 0.0001. Error bars represent SEMs. ns: not significant. IF: immunofluorescence microscopy, TEM: transmission electron microscopy, NS: negative staining method, CS: cross-sectioning method.

TEM analysis revealed multiple round to oval mitochondria-like structures with distinct inner and outer membranes, as shown in lower- and higher-resolution images in Fig. 7d–i. The size distribution graph for the TEM (negative staining) analysis of these experiments is shown in Fig. 7j, with an average size ranging from 39 nm to 997.5 nm. Next, the datasets from mitochondria-specific labeling (TOM20 IF) were compared with data from the same experiments obtained using label-free methods, including Videodrop and TEM (both negative staining and cross-sectioning). The results showed mean sizes of 333.9 ± 3.9 nm, 224.1 ± 5.7 nm, and 193.8 ± 6 nm, as detected by Videodrop, TEM-negative staining, and TEM-cross-sectioning, respectively. Additionally, mitochondria-like structures measured by cross-sectioning TEM had a mean size of 457.1 ± 2 nm (Fig. 7k). Statistically significant differences (*p* < 0.0001) were detected between the mean diameters measured via IF and/or mitochondria-like structures compared to those measured by Videodrop and TEM (negative staining or cross-sectioning). Interestingly, no significant differences were observed between the mean particle diameters measured by Videodrop and TEM average diameter using both methods (Fig. 7k). Although adjusting the thresholds for particle mean diameters above 80 nm or below 1000 nm (size detection limits of Videodrop) altered the mean diameters measured by each method (Supplementary Figure 2), it did not statistically affect the results.

Moreover, the automated quantification of TOM20+ particles revealed a mean particle concentration of 7.3 × 10⁸ ± 1 × 10⁸ part/mL (mean total count: 1 × 10⁸ ± 1.6 × 10⁷ particles), which was significantly lower compared to the concentration measured by Videodrop (8.8 × 10¹¹ ± 6 × 10¹⁰ particles/mL, mean total count: 1.3 × 10¹¹ ± 9.4 × 10⁹ particles) for the same experiments (n = 3). Protein concentration and total protein content for the same samples were 10.28 ± 0.57 mg/mL and 1.54 ± 0.08 mg, respectively. Notably, the mitochondria protein concentration measured by the Bradford assay showed less variability compared to Videodrop concentration, whereas particle concentration determined by TOM20 IF exhibited greater variability. Supplementary Table 2 summarizes the quantification data for the samples used in this assessment. Comparing size and density measurements between TOM20 IF and Videodrop suggests that the immunofluorescence method may overestimate the size of isolated mitochondria due to the presence of aggregated or overlapping structures (Fig. 7b–bi), while underestimating the overall particle concentration.

## Discussion

Mitochondrial transplantation offers a promising therapy for mitochondrial dysfunction by delivering freshly isolated mitochondria to target cells. However, quantifying mitochondrial density and size before transplantation is challenging due to their fragility post-isolation and susceptibility to freezing-thawing damage. Isolated mitochondria are highly sensitive to environmental changes, with the isolation process itself potentially disrupting their structure and function. McCully et al. reported that mitochondria remain stable at 4°C for up to 120 minutes, but beyond this, their functionality declines, highlighting their vulnerability and the need for rapid processing^8,44^. In this study, we demonstrated the effectiveness of label-free interferometry light microscopy, using Videodrop technology, for rapid and precise measurement of density and size in mitochondria freshly isolated from hMSCs. Its accuracy and reliability were evaluated through comparisons with widely used techniques.

Recently, there has been increasing interest in harnessing mitochondria derived from hMSCs as a biotherapeutic strategy^21,22^. This interest stems from the favorable properties of hMSCs, including their ease of availability, capacity for expansion in culture, and low immunogenicity^20^. These attributes position hMSCs as a promising source for allogeneic mitochondrial transplantation.

We first expanded hMSCs *in vitro* across 9 independent experiments before isolating mitochondria. Various techniques exist for mitochondrial isolation, including differential and density gradient centrifugation, differential filtration, immunomagnetic separation (anti-TOM20/anti-TOM22 functionalized magnetic beads^43^), subcellular fractionation with detergents, microfluidics, and commercial isolation kits (e.g., ThermoFisher^45^, Sigma-Aldrich^46^, Miltenyi Biotec or Abcam^47^). Here, we used the widely cited Abcam kit to isolate hMSC-derived mitochondria.

Isolated mitochondria were characterized using dual staining with anti-TOM20 (outer membrane marker)^48^ and MitoTracker Red CMXRos, a cationic dye that binds irreversibly to polarized, viable mitochondria^36,37^. These markers are well-established for mitochondrial characterization^23,27,49,50^. Western blotting further confirmed mitochondrial quality and integrity by detecting five OXPHOS complex subunits and six key structural proteins. Structural integrity was validated by electron microscopy. Given the size heterogeneity of mitochondria^51^, we analyzed the distribution of mitochondria-like structures in mitochondria-rich preparations using TEM cross-sections. The size ranged from 105 to 1,346 nm, with mean diameters of 354 nm (minimum), 489 nm (maximum), and 421 nm (average). Only 1.92% of particles exceeded 1 µm. Yao et al. (2022) reported a similar range (100–1,000 nm) for hMSC-derived mitochondria but did not provide detailed TEM data or quantification^52^. While TEM provides precise structural information about the diameter of isolated mitochondria, it is a time-consuming, costly, low throughput and complex method, making it not practical for the rapid characterization and quantification of freshly isolated mitochondria prior to transplantation.

Various alternative techniques have been used to characterize mitochondria in terms of density and size. While a hemocytometer can be a cost-effective method, its accuracy is highly debated, as it relies on the experimenter’s technique and is prone to personal errors^27^. Immunofluorescent staining with mitochondria membrane potential indicators (e.g., MitoTracker probes) or specific markers (e.g., anti-TOM20) is commonly used to assess mitochondria number^7,23,24^. Labeled mitochondria can be counted using a fluorescent microscope and analysis software. However, the main drawback of this method is that the mitochondria often aggregate and overlap, making it inaccurate and unreproducible^11^, as confirmed also by our measurements of TOM20+ mitochondria. Furthermore, size quantification via immunofluorescent staining is limited by the diffraction properties of the light. Indeed, the size of the fluorescent detected object matches the real size of the object only if its size is above the diffraction limit of the light (∼250nm). For smaller objects, which concerns the majority of the nanoparticles detected in mitochondria enriched samples, the size will be overestimated, which is consistent with our observations. Alternative methods like flow cytometry^5,26^ and nanoflow cytometry (NanoFCM)^25^ can quantify isolated mitochondria with immunofluorescence staining. While flow cytometry provides specific information on count and size, it is time-consuming and costly due to the need for sample labeling, limiting its utility for biotherapeutic applications in clinical settings.

Alternatively of the physical parameters quantification, a growing number of studies have used total protein measurements to indirectly estimate mitochondrial concentration^11^. Various protein assay techniques, including bicinchoninic acid (BCA), Bradford, Biuret, and Lowry methods, are commonly employed for this purpose^53–56^. These methods rely on a standard protein curve to compare the sample’s total protein content. In this study, we measured mitochondrial protein concentration and found strong correlation between Videodrop particle concentration and mitochondrial protein levels. However, the main limitation of these methods is their reliance on total protein content, which can vary based on mitochondrial purity. Purity is heavily influenced by the isolation method, experimental conditions, and user variability. Since total protein measurement requires mitochondrial lysis, these methods do not provide size (structural) information, which is a crucial mechanical characteristic for assessing mitochondrial integrity and purity. Non-viable mitochondria, previously frozen mitochondria, mitochondrial fractions (proteins, complexes I–V), mitochondrial DNA and RNA, and exogenous ATP or ADP have been shown to be ineffective in providing cytoprotection following mitochondrial transplantation^7,57,58^. Therefore, minimal yet effective quality control requirements for biomedical applications include both quantification and structural characterization. To meet this need, particle counters have been developed to directly measure the number and size of mitochondria following isolation.

Particle counters are automated tools that minimize user-induced errors, offering a reliable means to assess the size and count of isolated mitochondria. A type of particle counter has been used in a clinical trial involving mitochondrial transplantation^7^. For instance, McCully’s team utilized a Multisizer 4e Coulter Counter (Beckman-Coulter) to determine mitochondrial numbers^27,57,59^. Preble et al. demonstrated that counting mitochondria with Coulter counter surpassed the hemocytometer^27^. The Coulter principle, based on changes in impedance as particles pass through an orifice, enables measurement of particle volume without relying on particle characteristics like refractive index. Typically used in hematology to count and size blood cells, Coulter counters can measure particles ranging from 200 nm to 1600 µm. Thus, Coulter counters can overestimate the diameter and undercount the number of isolated mitochondria due to their minimum detection size of 200 nm. Moreover, Coulter counters require significant amount of electrolyte (5 mL minimum), limiting its use to hydrophilic particles in large sample containers like beakers or vials, which can lead to low particle concentration. Finally, the high running costs and expensive workflow make it less practical for routine use.

In this study, we used a Videodrop instrument, a light interferometry-based method, to characterize isolated mitochondria for biotherapy. The process involves pipetting 5-7 µL of the sample into a small well on a glass coverslip, followed by cleaning after measurement. Settings such as particle detection threshold, tracking length, viscosity, and minimum particle count can be adjusted based on particle properties (e.g., refractive index, density). The optical system records particle movement through short video blocks (100 frames at 140 fps) acquired successively until the minimum particle count is reached. The precision improves with the number of particles (>300nm for all our measurements). Image processing allow to retrieve the interferometric signal where submicrometric particles (>80nm) can be followed in time. All detected particles contribute to particle concentration, while the hydrodynamic radius is determined from tracked particles only. Videodrop is known to measure concentrations between 10^8 and 10^10 particles/mL, with a size range of 80 to 1000 nm for biological particles.

We assessed Videodrop’s reliability in measuring mitochondrial concentration and size across 9 experiments with 5-7 serial dilutions and triplicates. The optimal concentration range for hMSC-derived mitochondria (7.5×10^7 to 2.7×10^9 particles/mL) showed linearity with theoretical concentrations. Videodrop particle concentrations correlated with Bradford protein assays and NucleoCounter® cell counts, confirming its accuracy in quantifying mitochondrial concentration.

To enhance the characterization of particles in the mitochondria samples, we analyzed the size data from Videodrop alongside particle concentration measurements. The integrity and size distribution of isolated mitochondria can be influenced by factors such as isolation technique, cell type, and the cells’ physiological state before isolation^51^. Videodrop data showed an average size of 315 nm for isolated hMSC-derived mitochondria, consistent with the 380 nm mean diameter reported by McCully et al. using Coulter counter (which has a minimum detection limit of 200 nm)^27^. Yao et al. reported a size range of 100-1000 nm for adipose tissue-derived hMSC mitochondria measured by dynamic light scattering (DLS), which aligns with their TEM results^52^.

Our TEM images confirm the isolation of polydisperse double membrane hMSCs-mitochondria, indicating that the isolated mitochondria exhibit a range of sizes and do not display uniformity in size distribution. To gain insight into the particle diameter that ILM was capable of measuring, we conducted a series of TEM analyses. We measured the minimum, maximum, and average diameter of objects exceeding (or not) the 80 nm or 1000 nm size detection limits of Videodrop, in both TEM images of cross-sectioned samples (with higher resolution) and TEM images of negative staining (with lower resolution). Considering all particles, our results demonstrate that the diameter detected by ILM (Videodrop) aligns with the mean maximum particle diameters detected in TEM images of negative staining (276 nm) but not with other measured diameters (minimum or average diameters by negative staining or all diameters measured by cross-sectioning methods). Since electron microscopy provides images with very high resolution, this was expected, as smaller particles generated during mitochondria isolation could decrease the mean particle diameter compared to interferometric light microscopy. TEM analysis revealed that approximately 5-7% (negative staining) and 18-20% (cross-sectioning) of total particles had diameters smaller than 80 nm, while particles exceeding 1000 nm were much less common (∼0.4% for negative staining and ∼0.2% for cross-sectioning TEM methods). Interestingly, reanalyzing the TEM data—considering the Videodrop’s lower size detection limit (80 nm)—no significant differences were observed between the mean particle diameter measured by Videodrop and the maximum or average diameters detected by negative staining TEM. However, the Videodrop mean diameter was still significantly higher compared to the mean diameters measured by cross-sectioning TEM, even after the 80 nm cut-off. These results suggest that while ILM, as used in Videodrop, may not provide the same high resolution as TEM of cross-sectioned samples, it is still sufficiently accurate and reliable for detecting particle diameter, similar to TEM of negative staining. It has been reported that negative-stain images inherently possess higher contrast compared to cryo-electron microscopy (cryo-EM), which has also been widely employed to detect isolated mitochondria^58^. This could be partly due to the use of heavy metal stains in negative staining, which enhance the visibility of the particles or structures of interest. Additionally, different sample processing for TEM negative staining vs cross section may affect the size of the particles. Also, for ILM the quantification of the mitochondria hydrodynamic diameter is based on the Stokes-Einstein equation, which is valid for spherical object submitted to thermal agitation inducing Brownian motion. While there is nothing indicating that mitochondria may exhibit motion other than due to thermal agitation, mitochondria exhibit a wide range of shapes from spherical to elongated. Here, by using ILM we assume that the Stokes-Einstein equation is valid and therefore we supposed that the objects are spherical. For TEM image analysis, we considered both the min diameter and the max diameter by calculating the average diameter. For ILM, it is not sure whether the hydrodynamic diameter obtained is related to the max diameter, the min diameter or a size in between both diameters. However, as hydrodynamic diameter measured by ILM was close to the maximum diameter measured by negative staining TEM, the diffusive behavior of mitochondria may resemble to the Brownian motion of spherical particles of size corresponding to the max diameter. Additionally, ILM size measurement gives access to the hydrodynamic diameter, which is usually higher than the geometric radius for nanoparticles. Nevertheless, ILM measurement is the closest to the size obtained by TEM and highly less time consuming than the other techniques, ILM should still be considered as a potential method for assessing mitochondria’s size as a quality control tool.

In this paper, we conducted side-by-side experiments comparing mitochondria characterization using TOM20 IF microscopy, TEM, Bradford assay for protein measurement, and Videodrop (label-free method) on the same samples. We observed frequent aggregation and overlapping of TOM20+ mitochondria in confocal images. Our results showed a significant difference in the mean diameter of TOM20+ particles measured by HCS Studio Cell Analysis software (938 nm) compared to Videodrop (333 nm) and TEM (219 nm for negative staining, 193 nm for cross-sectioning, and 457 nm for mitochondria-like structures). This difference may be due to the light diffraction limit. According to TEM quantification the majority of the particles contained in the mitochondria enriched fraction are above the limit of diffraction of the light (∼250nm). Meaning that the size of the fluorescent objects is going to be higher than the real particle size, therefore inducing an overestimation of the size by confocal fluorescent microscopy technique. No significant differences were found between the mean diameters measured by Videodrop and TEM methods, with results remaining consistent even after introducing cut-offs. Bahnemann et al. reported a size range of 0.5–2.5 μm for CHO-K1-derived mitochondria isolated using Dounce or ultrasound homogenization methods^23^, which were labeled with the same anti-TOM20 antibody as in our study. These findings, altogether reinforced the advantage of Videodrop’s label-free interferometry for accurate size measurement.

Particle quantification data showed a lower concentration measured by the TOM20 immunofluorescence method compared to interferometry analysis by Videodrop. This discrepancy may stem from TOM20+ mitochondria structures typically being larger in mitochondria-rich samples. Also, the resolution of confocal microscopy does not allow to differentiate two close nanoparticles from one bigger. High-resolution TEM cross-sectioning revealed a mean size of 421 nm for mitochondria-like structures, which is significantly lower than that measured by TOM20 immunofluorescence, but aligns with the size observed in TEM images. These results suggest that conventional immunolabeling methods can lead to mitochondrial aggregation, hindering accurate quantification of isolated mitochondria compared to TEM or Videodrop methods.

Finally, it is important to note that, as a label-free method, the accuracy of Videodrop particle counter measurements for isolated mitochondria size or density largely depends on sample purification. which in turn is influenced by the cell/tissue source and isolation methods. Indeed, ILM does not allow to differentiate between mitochondria-like objects and other particles included in cell secretome. Whether the findings of this study apply to hMSCs-mitochondria purified using different techniques or derived from other cell sources, such as adipose-derived tissues, can only be confirmed through direct comparison.

## Conclusion

Overall, our findings, which include comparisons between ILM by Videodrop, protein assays by Bradford method, negative staining and cross-sectioned TEM, and TOM20 IF methods, demonstrate the reliability of Videodrop as an efficient device for quality control analysis of freshly isolated hMSCs-derived mitochondria, suitable for use in bioprocessing, engineering, production, or dosing for biotherapeutic applications. Additionally, Videodrop, to our knowledge, is the most time-efficient label-free particle counter, making it ideal for mitochondrial transplantation studies or biotherapies requiring rapid characterization and quantification of freshly isolated mitochondria before transplantation. The data presented in this study should have broad applications in the emerging field of mitochondria transplantation, particularly in translational studies that require quick and efficient characterization of small volumes (few microliters) of precious human mitochondria in preclinical, pharmaceutical, or clinical settings.

## Methods

### Culture of human adipose tissue-derived mesenchymal stem cells (hMSCs)

The mitochondria were isolated from commercially purchased human adipose tissue-derived mesenchymal stem cells (CellEasy batch number N°9297). The hMSCs were cultured under standard tissue culture condition, at 37 °C and 5% CO2. Cells were grown in T175 culture flasks using α-Minimal Essential Medium (α-MEM) supplemented with 10% fetal bovine serum (FBS). Twenty mL of media was used to ensure complete coverage of the T150 flask. Medium was changed every 3 days. MSCs were grown to 90% confluence before being passage. Cells were lifted from the flask first by aspirating the media and washing in their flasks with phosphate buffered saline (PBS, Gibco, Thermofisher). Washed cells were then incubated in 0.05% trypsin ethylenediaminetetraacetic acid (EDTA, Gibco, Thermofisher) for 2 to 3 minutes in an incubator at 37°C and 5% CO_2_. The flasks were tapped to help mechanical detachment of the cells. An equal volume of α-MEM with 10% FBS was added to neutralize the trypsin. The cell solution was then centrifuged at 300 g for 10 mins and the supernatant was discarded. Cells were quantified using the NucleoCounter® NC-200™ automated cell analyzer (NC-200, Chemometec) to determine the total cell counts and cell viability for each experiment. Only cells between passages 4 to 8 were used for mitochondria isolations.

### Isolation of mitochondria from hMSCs

Mitochondria were isolated from hMSCs using a commercial mitochondrial isolation kit for cultured cells (ab110170, Abcam) ^47,60,61^. Briefly, cells were pelleted by centrifugation at 300 g. Cells were quickly frozen and thawed to weaken the cell membranes, and suspend in Reagent A at 5.0 mg protein/ml (approximately 1ml per 25 million cells) for 10 minutes on ice. Cell disruption was carried out using a sonication probe (Sonicator FB50, Fisher Scientific). Ultrasound was applied through 3 cycles of 10-s bursts at 20% power and with 10-s intervals in between the bursts (30-s in total active phase). The cell suspensions were cooled on ice during sonication.

Then, the homogenate was centrifuged at 1,000 g for 10 minutes at 4°C to pellet the remaining cells, cell debris and nuclei. The supernatant (SN) #1 containing mitochondria was saved and the pellet was resuspended in Reagent B to the same volume as Reagent A. The cell rupturing step was repeated to release mitochondria from the remaining intact cells. Same steps were applied and the supernatant SN #2 was saved and the pellet was discarded. SNs #1 & #2 were combined, thoroughly mixed and centrifuged again at 12,000g for 15 minutes at 4°C. After that step, the pellet was saved as an isolated mitochondria mass. The pellet was washed with 200 µL of Reagent C supplemented with Protease Inhibitors (11697498001, Roche). Then, it was centrifuged again (12,000g for 15 minutes at 4°C), and then aliquoted for characterization on fresh isolated mitochondria or frozen and store at −80°C until the mitochondrial quality assays described below were performed.

### Visualization of isolated mitochondria

Mitochondria viability was assessed using MitoTracker Red CMXROS (M7512, Invitrogen) staining and observed using a transmission microscope equipped with filter cubes (EVOS M5000, ThermoFisher). MitoTracker Red CMXROS is a red-fluorescent dye that stains intact mitochondria and its accumulation is dependent upon mitochondria membrane potential. Freshly isolated mitochondria were stained with MitoTracker Red (200 nM) for 40 minutes, washed twice with PBS 1X, resuspended in 10 µL of PBS, placed on slides and covered for microscopic observation using EVOS M5000 Imaging System (ThermoFisher).

### Immunofluorescence imaging of isolated mitochondria

Freshly Isolated mitochondria were characterized by mitochondria specific immunostaining. Samples of isolated mitochondria were incubated for 30 min with a primary rabbit anti-mouse TOM20 (translocase of outer mitochondrial membrane 20 homolog) antibody (Proteintech), which binds specifically to the TOM20 complex located in the mitochondrial outer membrane. After incubation, samples were centrifuged (12000 g, 15 min, 4°C) and washed twice with PBS 1X. Then, the mitochondria were incubated (30 min, in dark) with a secondary antibody (Goat anti-Rabbit IgG), which was conjugated with green fluorescent AlexaFluor 488 (A-11008 Invitrogen). Visualization was achieved by a Cell Insight CX7-LZR automated confocal spinning-disk fluorescence microscope (ThermoFisher Scientific). Images were captured and analyzed simultaneously to their acquisition thanks to the HCS Studio Cell Analysis software package capable of analyzing images.

### Mitochondrial yield for quantitation of proteins

The concentration of isolated raw mitochondrial protein was determined by Bradford protein assay^62,63^ according to the manufacturer’s protocol against a standard calibration curve constructed using BSA. Bradford assay was performed in 96-well plates using Bradford Reagent (B6916, Sigma Aldrich). The plate was read at 595 nm absorbance detection by a Microplate Spectrophotometer (Molecular Devices, SpectraMax iD3 plate reader). Mitochondrial protein yields were calculated in milligrams per mL based on the measured signal intensities and deduced from the calibration curve.

### Western blot analysis of mitochondria protein markers

Mitochondria obtained from hMSCs were homogenized and lysed in ice-cold 30 mM Tris-EDTA, pH 7.2, containing 1 mM DTT, 1% (v/v) Triton X-100, 10% (w/v) anti-phosphatase cocktail and 14% (w/v) anti-protease cocktail (11697498001, Roche), for 30 min on ice. After that, the homogenate was gently mixed on ice for 2 minutes on agitating plate. The proteins in the supernatants were then quantified using a Bradford assay and used for Western blot analysis. Twenty µg of total protein of each sample was loaded and separated on 4–20% Mini-PROTEAN® TGX™ Precast Protein Gels (Bio-Rad) and electroblotted onto membranes of Trans-Blot Turbo Midi 0.2 µm PVDF Transfer Packs (Bio-Rad). After being rinsed in TBS-Tween 20 buffer (TBST), the membranes were blocked for 1 h in the blocking buffer (EveryBlot Blokcking Buffer, BioRad), then washed with TBST and incubated overnight at 4°C with either (i) rabbit anti-TOM20 (1:5000, 11802-1-AP, ThermoFisher) primary antibody; (ii) total oxidative phosphorylation (OXPHOS) Human WB Antibody Cocktail (MS601) (1:700, ab110411, Abcam) that contains 5 antibodies, one each against Complex I subunit NDUFB8 (NADH dehydrogenase (ubiquinone) 1 beta subcomplex subunit 8), Complex II subunit, Complex III subunit Core 2, Complex IV subunit II, and Complex V ATP synthase subunit alpha as an optimized premixed cocktail (which are the subunits that are labile when their complexes are not assembled); (iii) Membrane Integrity WB Antibody Cocktail (MS620) (1:250, ab110414, Abcam) that contains 5 antibodies against representative proteins for each of the primary structures of the mitochondrion: Human VDAC1 (voltage-dependent anion channel) or Porin located in the outer membrane mitochondrial protein; Cyclophilin F located in the mitochondrion matrix; Ubiquinol-Cytochrome C Reductase Core Protein I, which is a part of the mitochondrial electron transport chain located in the inner mitochondrial membrane; Cytochrome C located in the mitochondrial intermembrane/intercristae spaces; and ATP5A located in the inner mitochondria membrane and matrix (to evaluate the structural integrity of isolated mitochondria) or (iv) rabbit anti-GAPDH antibody as a loading control antibody (1: 2500, ab9485, Abcam). After washing, the blots were incubated with the corresponding goat anti-rabbit (1:2000, 7074 Cell Signaling) or goat anti-mouse (1:2000, 7076, Cell Signaling) horseradish peroxidase (HRP)-conjugated secondary antibodies for 2h at RT. Peroxidase activity was revealed with a chemiluminescent detection kit (ECL Plus substrate, GE Healthcare). Protein expression visualization and acquisitions were performed by Syngene PXi (UK). Documentation was accomplished using GeneSys image capture software (UK). Post-spin supernatant was loaded as control at 20 μg for each experiment and immunoblotting.

### Interferometric Light Microscopy measurements of isolated mitochondria using Videodrop

The average hydrodynamic radius and the concentration of isolated mitochondria samples from 9 experiments were determined by ILM using Videodrop technology (Myriade, Paris, France). Samples were diluted in PBS 1X at 5-7 serial dilutions (d1, d50, d100, d200, d400, d800, d1600). Each dilution was measured in triplicates. Signals from the PBS 1X were checked and always found under the detection limit. For the measurement, drops of 7 µL each were loaded on a round-shaped cover slip using a micropipette before lifting the stage towards the objective. The software QVIR 2.6.0 (Myriade, Paris, France) was then used to illuminate the sample using a 2 W blue LED light. The images were recorded by a complementary metal-oxide-semiconductor high-resolution high-speed camera. Each particle scatters the incident light and behave like a secondary source of light. From the interferences of these secondary light sources, an interference pattern is generated by the software. This allows sub-micrometric particles to be detected quickly and followed in time. The concentration of the particles corresponds to the number of detected particles within the measured volume, which depends on the microscope characteristics and the particles’ size. The hydrodynamic radius was estimated by tracking the position of the imaged particle within the recorded movie. Indeed, the particles are moving due to thermal agitation. As passive particles in a pure viscous medium, they exhibit a Brownian motion, where the mean square displacement follows this relation:

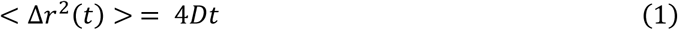

*D* is the diffusion coefficient and *t* the time. The diffusion coefficient of each particle is deduced from its mean square displacement, itself deduced from the particle trajectory. Then, the hydrodynamic radius (*R*_*h*_) (= half of the hydrodynamic diameter) is calculated using the Stokes-Einstein equation (Eq. 1), where *D* is the diffusion coefficient, *k*_*B*_ the Boltzmann constant, *T* the temperature (measured for each Videodrop measurement), *η* the dynamic viscosity (usually the one of water):

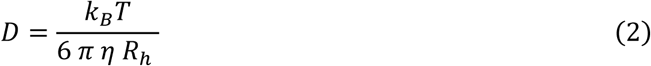

Particles with smaller masses move within a larger volume than particles with larger masses.

The minimum particles to be tracked were set at 300 particles. The maximum number of videos was set at 50 and the number of frames was 100. Minimum saturation was considered at 92%. In accordance with the manufacturer’s specifications, a relative threshold of 4.2 was applied for detection. The consideration of macroparticles was enabled using a minimum radius of 10 and a minimum number of hot pixels of 150, as well as drift compensation. Concerning the tracking settings, a maximum of two jumps was tolerated for a minimal length track of 10 frames. The doublet detector (QVIR software version 2.6.0.7378) was used for all the measurements. After measurements, the majority of the results were saved both as a PDF report and as a CSV file.

### Electron Microscopy

***Negative-staining method:*** Isolated mitochondria (5 µL) were directly adsorbed onto a carbon film membrane on a 300-mesh copper grid, stained with 1% uranyl acetate dissolved in distilled water, and dried at RT. Grids were examined using a Hitachi HT7700 electron microscope operated at 80 kV (Elexience) and the images acquired with a charge coupled device camera (AMT).

***Cross-sectioned method:*** Suspensions (45 μL) of mitochondria were pelleted at 12,000 ×g for 15 min, the supernatant was carefully removed, and the pellets were fixed with 2% glutaraldehyde in 0.1 M Na cacodylate buffer pH 7.2, for 1 h at RT. The supernatant was removed and the pellets were washed 3 times in PBS 1X. Samples were then contrasted with Oolong Tea Extract (OTE) 0.2% in cacodylate buffer, postfixed with 1% osmium tetroxide containing 1.5% potassium cyanoferrate for 1 h. Following three additional PBS 1X washes, the pellet was dehydrated through a graded series (30% to 100%) and substituted gradually in mix of ethanol-epon and embedded in 100% Epon (Delta microscopie, France) over 24 h. The pellet was embedded in a final change of Epon, cured at 37 °C overnight, followed by additional hardening at 65 °C for two more days. Ultrathin sections (80 nm) were collected onto 200 mesh copper grids, stained with 2% uranyl acetate in 50% methanol for 10 min, and counterstained with 1% lead citrate for 7 min. Grids were examined with Hitachi HT7700 electron microscope operated at 80kV (Milexia – France), and images were acquired with a charge-coupled device camera (AMT). This work was carried out at, and with the expertise of, MIMA2 MET – GABI, INRA, Agroparistech, 78352 Jouy-en-Josas, France.

***Scanning Electron Microscopy (SEM):*** Samples were mounted on aluminum stubs (32 mm diameter) with carbon adhesive discs (Agar Scientific, Oxford Instruments SAS, Gomez-la-Ville, France) and visualized by field emission gun scanning electron microscopy (SEM FEG) as secondary-electron images (2 keV, spot size 30) under high-vacuum conditions with a Hitachi SU5000 instrument (Milexia, Saint-Aubin, France). SEM analyses were performed at the Microscopy and Imaging Platform MIMA2 (INRAE, Jouy-en-Josas, France) DOI: MIMA2, INRAE, 2018. Microscopy and Imaging Facility for Microbes, Animals and Foods.

### Statistics and reproducibility

Statistical analyses were performed with Prism v.8.0.2 software (GraphPad Software, LLC) using unpaired two-tailed Mann–Whitney test (non-parametric test) and ordinary one-way or two-way analysis of variance (ANOVA). The notations for the different levels of significance are indicated as follows: *p < 0.05, **p < 0.01, ***p < 0.001, ****p < 0.0001. For all the experiments, a p-value inferior to 0.05 is considered significant. Data are presented as means + SEM. Number of experiments or replicates used per experiment have been mentioned in the results section as well as in the figure legends.

## Author contribution

Conceptualization, S.M., and A.S.B.; data curation, S.M. and K.A.; formal analysis, S.M. and K.A.; funding acquisition, A.S.B. and F.G. and; investigation, S.M. and K.A.; methodology, S.M., C.R., D.A., M.B. and K.A.; project administration, S.M. and A.S.B.; resources, F.G., A.S.B.; software, S.M., K.A. and A.S.B.; supervision, S.M., A.S.B. and KA; validation, S.M., K.A., M.B., F.G., A.S.B.; visualization, S.M.; writing—original draft, S.M.; writing—review and editing, S.M., K.A., M.B., F.G. and A.S.B. All authors have read and agreed to the published version of the manuscript.

## Acknowledgements

The authors would like to acknowledge Marie Berger (Myriade Lab) for sharing her deep knowledge about interferometric light microscopy technology and for her availability for all the discussions about this work. Videodrop and CellInsight CX7-LZR are part of the IVETh expertise facility. IVETh was supported by the IdEx Université Paris Cité, ANR-18-IDEX-0001 (IVETh plateform), by the Region Ile de France under the convention SESAME 2019 - IVETh (EX047011) and by the Region Ile de France and Banque pour l’Investissement (BPI) under the convention Accompagnement et transformation des filières projet de recherche et développement N° DOS0154423/00 & DOS0154424/00, DOS0154426/00 & DOS0154427/00, Agence Nationale de la Recherche through the program France 2030 “Integrateur biotherapie-bioproduction” (ANR-22-AIBB-0002). IVETh is also supported by Region Île-de-France in the framework of DIM BioConvS. This project has received funding from CNRS and from the European Research Council (ERC) under the European Union’s Horizon 2020 research and innovation program (grant agreement No.852791). We would like also to acknowledge technical assistance from the Microscopy and Imaging Platform MIMA2 (INRAE, Jouy-en-Josas, France) DOI: MIMA2, INRAE, 2018; Microscopy and Imaging Facility for Microbes, Animals and Foods. In particular, we would like to thank Christine Longin and Vlad Costache from the INRAE MIMA2 platform for their valuable help in performing TEM and SEM analyses. The authors declare that artificial intelligence is not used in this study.

## References

1 Norat, P. et al. Mitochondrial dysfunction in neurological disorders: Exploring mitochondrial transplantation. NPJ Regen Med 5, 22, doi:10.1038/s41536-020-00107-x (2020).

2 Roushandeh, A. M., Kuwahara, Y. & Roudkenar, M. H. Mitochondrial transplantation as a potential and novel master key for treatment of various incurable diseases. Cytotechnology 71, 647–663, doi:10.1007/s10616-019-00302-9 (2019).

3 Rath, E., Moschetta, A. & Haller, D. Mitochondrial function - gatekeeper of intestinal epithelial cell homeostasis. Nat Rev Gastroenterol Hepatol 15, 497–516, doi:10.1038/s41575-018-0021-x (2018).

4 Liu, D. et al. Intercellular mitochondrial transfer as a means of tissue revitalization. Signal Transduct Target Ther 6, 65, doi:10.1038/s41392-020-00440-z (2021).

5 Al Amir Dache, Z., et al. Blood contains circulating cell-free respiratory competent mitochondria. FASEB J 34, 3616–3630, doi:10.1096/fj.201901917RR (2020).

6 Lightowlers, R. N., Chrzanowska-Lightowlers, Z. M. & Russell, O. M. Mitochondrial transplantation-a possible therapeutic for mitochondrial dysfunction?: Mitochondrial transfer is a potential cure for many diseases but proof of efficacy and safety is still lacking. EMBO Rep 21, e50964, doi:10.15252/embr.202050964 (2020).

7 McCully, J. D. et al. Injection of isolated mitochondria during early reperfusion for cardioprotection. Am J Physiol Heart Circ Physiol 296, H94–H105, doi:10.1152/ajpheart.00567.2008 (2009).

8 McCully, J., del Nido, P. J. & Emani, S. M. Therapeutic mitochondrial transplantation. Current Opinion in Physiology 27, doi:ARTN 100558 10.1016/j.cophys.2022.100558 (2022).

9 Gomzikova, M. O., James, V. & Rizvanov, A. A. Mitochondria Donation by Mesenchymal Stem Cells: Current Understanding and Mitochondria Transplantation Strategies. Front Cell Dev Biol 9, 653322, doi:10.3389/fcell.2021.653322 (2021).

10 Mitochondria transplantation, list of results. ClinicalTrials.gov, <https://clinicaltrials.gov/search?intr=%22mitochondria%22%20transplantation&viewType=Table&page=2> (

11 Kubat, G. B., Ulger, O. & Akin, S. Requirements for successful mitochondrial transplantation. J Biochem Mol Toxicol 35, e22898, doi:10.1002/jbt.22898 (2021).

12 Mozafari, S., Peruzzotti-Jametti, L. & Pluchino, S. Mitochondria transfer for myelin repair. J Cereb Blood Flow Metab, 271678X251325805, doi:10.1177/0271678X251325805 (2025).

13 Han, D. et al. Mesenchymal Stem/Stromal Cell-Mediated Mitochondrial Transfer and the Therapeutic Potential in Treatment of Neurological Diseases. Stem Cells Int 2020, 8838046, doi:10.1155/2020/8838046 (2020).

14 Loussouarn, C., Pers, Y. M., Bony, C., Jorgensen, C. & Noel, D. Mesenchymal Stromal Cell-Derived Extracellular Vesicles Regulate the Mitochondrial Metabolism via Transfer of miRNAs. Front Immunol 12, 623973, doi:10.3389/fimmu.2021.623973 (2021).

15 Morrison, T. J. et al. Mesenchymal Stromal Cells Modulate Macrophages in Clinically Relevant Lung Injury Models by Extracellular Vesicle Mitochondrial Transfer. Am J Respir Crit Care Med 196, 1275–1286, doi:10.1164/rccm.201701-0170OC (2017).

16 Dutra Silva, J., et al. Mesenchymal stromal cell extracellular vesicles rescue mitochondrial dysfunction and improve barrier integrity in clinically relevant models of ARDS. Eur Respir J 58, doi:10.1183/13993003.02978-2020 (2021).

17 Thomas, M. A. et al. Human mesenchymal stromal cells release functional mitochondria in extracellular vesicles. Front Bioeng Biotechnol 10, 870193, doi:10.3389/fbioe.2022.870193 (2022).

18 Wang, C. et al. Postischemic Neuroprotection Associated With Anti-Inflammatory Effects by Mesenchymal Stromal Cell-Derived Small Extracellular Vesicles in Aged Mice. Stroke 53, e14–e18, doi:10.1161/STROKEAHA.121.035821 (2022).

19 Harman, R. M. et al. Single-cell RNA sequencing of equine mesenchymal stromal cells from primary donor-matched tissue sources reveals functional heterogeneity in immune modulation and cell motility. Stem Cell Res Ther 11, 524, doi:10.1186/s13287-020-02043-5 (2020).

20 Huaman, O. et al. Immunomodulatory and immunogenic properties of mesenchymal stem cells derived from bovine fetal bone marrow and adipose tissue. Res Vet Sci 124, 212–222, doi:10.1016/j.rvsc.2019.03.017 (2019).

21 Tan, Y. L. et al. Mesenchymal Stromal Cell Mitochondrial Transfer as a Cell Rescue Strategy in Regenerative Medicine: A Review of Evidence in Preclinical Models. Stem Cells Transl Med 11, 814–827, doi:10.1093/stcltm/szac044 (2022).

22 Mohammadalipour, A., Dumbali, S. P. & Wenzel, P. L. Mitochondrial Transfer and Regulators of Mesenchymal Stromal Cell Function and Therapeutic Efficacy. Front Cell Dev Biol 8, 603292, doi:10.3389/fcell.2020.603292 (2020).

23 Bahnemann, J. et al. In search of an effective cell disruption method to isolate intact mitochondria from Chinese hamster ovary cells. Engineering in Life Sciences 14, 161–169, 10.1002/elsc.201200182 (2014).

24 Ulger, O. et al. The effects of mitochondrial transplantation in acetaminophen-induced liver toxicity in rats. Life Sci 279, 119669, doi:10.1016/j.lfs.2021.119669 (2021).

25 Peruzzotti-Jametti, L. et al. Neural stem cells traffic functional mitochondria via extracellular vesicles. PLoS Biol 19, e3001166, doi:10.1371/journal.pbio.3001166 (2021).

26 Kubat, G. B. et al. The effects of mesenchymal stem cell mitochondrial transplantation on doxorubicin-mediated nephrotoxicity in rats. J Biochem Mol Toxicol 35, e22612, doi:10.1002/jbt.22612 (2021).

27 Preble, J. M. et al. Rapid isolation and purification of mitochondria for transplantation by tissue dissociation and differential filtration. J Vis Exp, e51682, doi:10.3791/51682 (2014).

28 Turkki, V., Alppila, E., Yla-Herttuala, S. & Lesch, H. P. Experimental Evaluation of an Interferometric Light Microscopy Particle Counter for Titering and Characterization of Virus Preparations. Viruses 13, doi:10.3390/v13050939 (2021).

29 Roose-Amsaleg, C. et al. Utilization of interferometric light microscopy for the rapid analysis of virus abundance in a river. Research in Microbiology 168, 413–418, doi:10.1016/j.resmic.2017.02.004 (2017).

30 Romolo, A. et al. Assessment of Small Cellular Particles from Four Different Natural Sources and Liposomes by Interferometric Light Microscopy. Int J Mol Sci 23, doi:10.3390/ijms232415801 (2022).

31 Sausset, R. et al. Comparison of interferometric light microscopy with nanoparticle tracking analysis for the study of extracellular vesicles and bacteriophages. bioRxiv, 2022.2010.2007.511248, doi:10.1101/2022.10.07.511248 (2022).

32 Lapras, B. et al. Real-time monitoring by interferometric light microscopy of phage suspensions for personalised phage therapy. Sci Rep 14, 31629, doi:10.1038/s41598-024-79478-w (2024).

33 Sabbagh, Q. et al. The von Willebrand factor stamps plasmatic extracellular vesicles from glioblastoma patients. Sci Rep 11, 22792, doi:10.1038/s41598-021-02254-7 (2021).

34 Richard, M. et al. Monitoring concentration and lipid signature of plasma extracellular vesicles from HR(+) metastatic breast cancer patients under CDK4/6 inhibitors treatment. J Extracell Biol 3, e70013, doi:10.1002/jex2.70013 (2024).

35 Alexandre, L. et al. Investigating Extracellular Vesicles in Viscous Formulations: Interplay of Nanoparticle Tracking and Nanorheology via Interferometric Light Microscopy. Small Science 5, 2400319, 10.1002/smsc.202400319 (2025).

36 Pendergrass, W., Wolf, N. & Poot, M. Efficacy of MitoTracker Green and CMXrosamine to measure changes in mitochondrial membrane potentials in living cells and tissues. Cytometry A 61, 162–169, doi:10.1002/cyto.a.20033 (2004).

37 Marcondes, N. A. et al. Comparison of JC-1 and MitoTracker probes for mitochondrial viability assessment in stored canine platelet concentrates: A flow cytometry study. Cytometry A 95, 214–218, doi:10.1002/cyto.a.23567 (2019).

38 Boccara, M. et al. Full-field interferometry for counting and differentiating aquatic biotic nanoparticles: from laboratory to Tara Oceans. Biomedical Optics Express 7, 3736–3746, doi:10.1364/Boe.7.003736 (2016).

39 Pickles, S., Arbour, N. & Vande Velde, C. Immunodetection of outer membrane proteins by flow cytometry of isolated mitochondria. J Vis Exp, 51887, doi:10.3791/51887 (2014).

40 Yamamoto, H. et al. Dual role of the receptor Tom20 in specificity and efficiency of protein import into mitochondria. Proc Natl Acad Sci U S A 108, 91–96, doi:10.1073/pnas.1014918108 (2011).

41 Franko, A. et al. Efficient isolation of pure and functional mitochondria from mouse tissues using automated tissue disruption and enrichment with anti-TOM22 magnetic beads. PLoS One 8, e82392, doi:10.1371/journal.pone.0082392 (2013).

42 McCully, J. D., Cowan, D. B., Emani, S. M. & Del Nido, P. J. Mitochondrial transplantation: From animal models to clinical use in humans. Mitochondrion 34, 127–134, doi:10.1016/j.mito.2017.03.004 (2017).

43 Rambold, A. S., Kostelecky, B., Elia, N. & Lippincott-Schwartz, J. Tubular network formation protects mitochondria from autophagosomal degradation during nutrient starvation. Proc Natl Acad Sci U S A 108, 10190–10195, doi:10.1073/pnas.1107402108 (2011).

44 Cloonan, S. M. et al. Mitochondrial iron chelation ameliorates cigarette smoke-induced bronchitis and emphysema in mice. Nat Med 22, 163–174, doi:10.1038/nm.4021 (2016).

45 Morioka, E. et al. Mitochondrial LETM1 drives ionic and molecular clock rhythms in circadian pacemaker neurons. Cell Rep 39, 110787, doi:10.1016/j.celrep.2022.110787 (2022).

46 Yano, M. et al. Functional analysis of human mitochondrial receptor Tom20 for protein import into mitochondria. J Biol Chem 273, 26844–26851, doi:10.1074/jbc.273.41.26844 (1998).

47 Chazotte, B. Labeling mitochondria with MitoTracker dyes. Cold Spring Harb Protoc 2011, 990–992, doi:10.1101/pdb.prot5648 (2011).

48 Cottet-Rousselle, C., Ronot, X., Leverve, X. & Mayol, J. F. Cytometric assessment of mitochondria using fluorescent probes. Cytometry A 79, 405–425, doi:10.1002/cyto.a.21061 (2011).

49 Woods, D. C. Mitochondrial Heterogeneity: Evaluating Mitochondrial Subpopulation Dynamics in Stem Cells. Stem Cells Int 2017, 7068567, doi:10.1155/2017/7068567 (2017).

50 Yao, X. et al. In-cytoplasm mitochondrial transplantation for mesenchymal stem cells engineering and tissue regeneration. Bioeng Transl Med 7, e10250, doi:10.1002/btm2.10250 (2022).

51 Azimzadeh, P. et al. Comparison of three methods for mitochondria isolation from the human liver cell line (HepG2). Gastroenterol Hepatol Bed Bench 9, 105–113 (2016).

52 Ko, S. F. et al. Hepatic (31) P-magnetic resonance spectroscopy identified the impact of melatonin-pretreated mitochondria in acute liver ischaemia-reperfusion injury. J Cell Mol Med 24, 10088–10099, doi:10.1111/jcmm.15617 (2020).

53 Lee, J. M. et al. Mitochondrial Transplantation Modulates Inflammation and Apoptosis, Alleviating Tendinopathy Both In Vivo and In Vitro. Antioxidants (Basel) 10, doi:10.3390/antiox10050696 (2021).

54 Huang, P. J. et al. Transferring Xenogenic Mitochondria Provides Neural Protection Against Ischemic Stress in Ischemic Rat Brains. Cell Transplant 25, 913–927, doi:10.3727/096368915X689785 (2016).

55 Preble, J. M., Kondo, H., Levitsky, S. & McCully, J. D. Quality control parameters for mitochondria transplant in cardiac tissue. JSM Biochem Mol Biol 2, 1008 (2014).

56 Masuzawa, A. et al. Transplantation of autologously derived mitochondria protects the heart from ischemia-reperfusion injury. Am J Physiol Heart Circ Physiol 304, H966–982, doi:10.1152/ajpheart.00883.2012 (2013).

57 Cowan, D. B. et al. Transit and integration of extracellular mitochondria in human heart cells. Sci Rep 7, 17450, doi:10.1038/s41598-017-17813-0 (2017).

58 Tharichen, L., Englmeier, R. & Forster, F. Sample Preparation of Isolated Mitochondria for Cryoelectron Tomography and In Situ Studies of Translation. Methods Mol Biol 2661, 75–88, doi:10.1007/978-1-0716-3171-3_5 (2023).

59 Feng, Y. et al. COX7A1 enhances the sensitivity of human NSCLC cells to cystine deprivation-induced ferroptosis via regulating mitochondrial metabolism. Cell Death Dis 13, 988, doi:10.1038/s41419-022-05430-3 (2022).

60 Toda, T. et al. Extremely low-frequency pulses of faint magnetic field induce mitophagy to rejuvenate mitochondria. Commun Biol 5, 453, doi:10.1038/s42003-022-03389-7 (2022).

61 Dong, L. et al. Mannose ameliorates experimental colitis by protecting intestinal barrier integrity. Nat Commun 13, 4804, doi:10.1038/s41467-022-32505-8 (2022).

62 Garcia-Cazarin, M. L., Snider, N. N. & Andrade, F. H. Mitochondrial isolation from skeletal muscle. J Vis Exp, doi:10.3791/2452 (2011).

63 Wieckowski, M. R. & Wojtczak, L. Isolation of crude mitochondrial fraction from cells. Methods Mol Biol 1241, 1–8, doi:10.1007/978-1-4939-1875-1_1 (2015).

