## Supplementary information for "Rapid and Efficient Quality Control Analysis of Isolated Mitochondria by Interferometric Light Microscopy"

* corresponding authors:

**Supplementary Table 1.** Size measurement of mitochondria-like structures detected by TEM (cross-sectioning method). Data presented considering particle diameter measurements from four independent experiments.

| Isolated mitochondria size | Maximum size | Minimum size | Average (maximum and minimum sizes) |
|---|---|---|---|
| **Range (nm)** | 116 – 1700 | 92 – 1049 | 105 – 1346 |
| **Mean (nm)** | 471.9 ± 44 | 344.4 ± 26 | 408.2 ± 35 |
| **>1 µm %** | 5.29 | 0.72 | 1.92 |

**Supplementary Table 2**. TOM20 IF method used for quantification of mitochondria samples showes higher particle diameter but lower particle concentration compared to Videodrop measurements.

| | **Quantification methods** | | |
|---|---|---|---|
| | **TOM20 IF** | **Videodrop** | **Protein Con.** |
| **Exp.** | Particles/mL | | mg/mL |
| i | 8,82E+08 | 9,11E+11 | 9,46 |
| ii | 8,01E+08 | 7,69E+11 | 9,98 |
| iii | 5,34E+08 | 9,79E+11 | 11,40 |
| **Mean** | **7,39E+08** | **8,86E+11** | **10,28** |
| **Exp.** | **Particle count** | | **Total protein (mg)** |
| i | 1,32E+08 | 1,37E+11 | 1,42 |
| ii | 1,20E+08 | 1,15E+11 | 1,50 |
| iii | 8,00E+07 | 1,47E+11 | 1,71 |
| **Mean** | **1,11E+08** | **1,33E+11** | **1,54** |

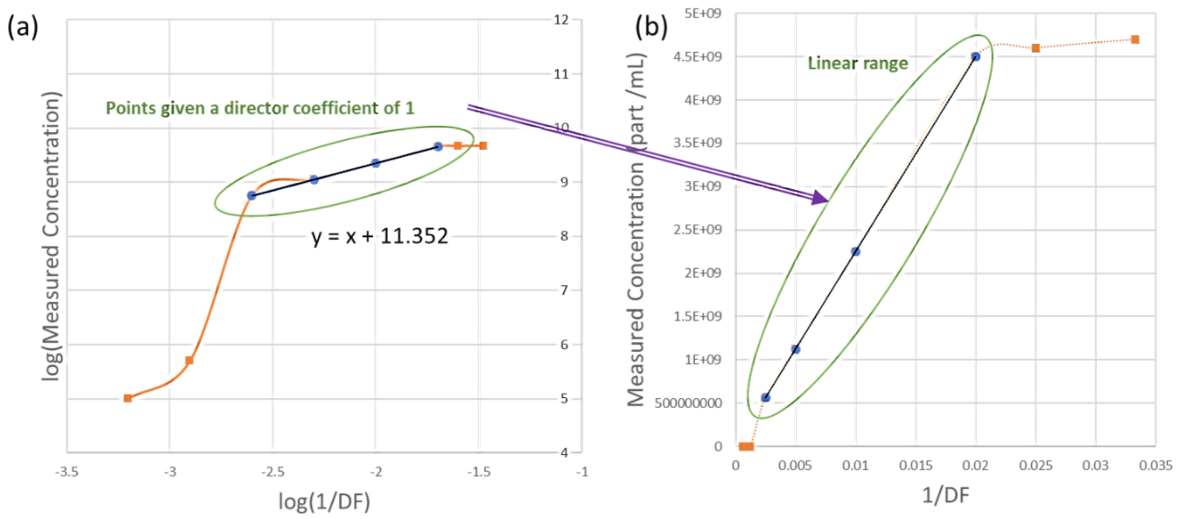


**Supplementary Figure 1.** Schematic illustration of the method V2 of determination of the theoretical concentration. (a) First, are identified the measurements for which a director coefficient of a=1 is obtained for a linear fit (y=ax+b) when plotting log(measured concentration) as a function of log(1/dilution factor [DF]). (b) Than, a verification is performed on the plot of the measured concentration as a function of 1/DF, to check for the linearity of the identified points. The theoretical concentration is then calculated based on the average value given by the identified points only. This method was applied independently for each of the 9 independent experiments.

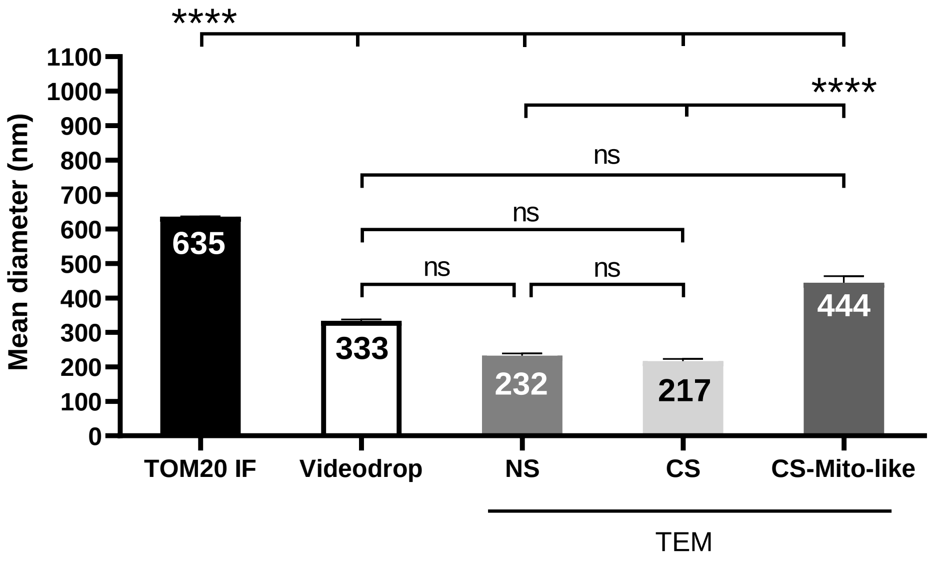


**Supplementary Figure 2.** Excluding particles with diameters below 80nm and above 1000nm (the detection limit of Videodrop) do not change the differences compared to including all particles (Fig. 7k). One-way ANOVA followed by Tukey's multiple comparison tests (n=3 experiments). **** $p < 0.0001$. Error bars represent SEMs. ns: non-significant, IF: immunofluorescence microscopy, TEM: transmission electron microscopy, NS: negative staining method, CS: cross-sectioning method.
